# A highly efficient gene disruption strategy reveals lipid co-regulatory networks

**DOI:** 10.1101/2020.11.24.395632

**Authors:** Takeshi Harayama, Tomomi Hashidate-Yoshida, Lucile Fleuriot, Auxiliadora Aguilera-Romero, Fumie Hamano, Keiken Ri, Ryo Morimoto, Delphine Debayle, Takao Shimizu, Howard Riezman

## Abstract

Gene disruption has been dramatically facilitated by genome editing tools. Despite improvements in gene disruption rates in cultured cells, clone isolation remains routinely performed to obtain mutants, potentially leading to artifacts due to clonal variation in cellular phenotypes. Here we report GENF, a highly efficient strategy to disrupt genes without isolating clones, which can be multiplexed. Using it, we obtained reliable lipidomics datasets from mutant cells without being affected by variances related to clone isolation. Through this, we found that an enzyme involved in congenital generalized lipodystrophy regulates glycerophospholipids with specific acyl-chains. We also demonstrate the possibility to dissect complex lipid co-regulatory mechanisms, explaining cell adaptations to altered lipid metabolism. With its simplicity and the avoidance of cloning-related artifacts, GENF is likely to contribute to many cell biology studies, especially those involving -omics approaches.

## INTRODUCTION

Gene disruption is a common strategy used in most biological fields to study the resulting phenotypic changes or to manipulate cellular functions. For this, it is obviously desirable that no phenotypic changes happen randomly during the process of gene disruption. Technologies based on the prokaryotic immune system CRISPR (clustered regularly interspaced palindromic repeats) have made gene disruption extremely easy.^1^ In the commonly used CRISPR-Cas9 system, the RNA-guided Cas9 nuclease is expressed in cells to induce double-strand breaks in genomic regions determined by an engineered single guide RNA (sgRNA), eventually leading to the formation of gene disruptive indels. The sequence of the sgRNA not only determines target specificity, but also affects the efficiency of cleavage.^2^ The efficiency of gene disruption by CRISPR-Cas9 has improved through advances in sgRNA design algorithms and delivery strategies of the components.^2–4^ However, complete gene disruption in the whole population of cells remains difficult, and therefore isolation of cellular clones after delivery of CRISPR-Cas9 component is a common practice to obtain mutated cells.^5^ However, it is known that clones of cell lines can have phenotypic variabilities.^6,7^ Thus, clone isolation could lead to accidental changes in cellular phenotypes irrespectively of the mutations induced by CRISPR-Cas9. This pitfall is starting to get attention in the literature, where the difficulty in finding correct phenotypes in mutant clones is discussed.^8,9^ This is problematic in most fields, for example in -omics studies where many variables are analyzed. Due to the presence of random variations introduced during clone isolation, it can be difficult to discriminate whether changes in -omics datasets are due to the genetic perturbation or due to clonal differences. In addition, clonal differences add noise to datasets, making quantitative changes between various mutants difficult to assess, and limiting the use of CRISPR-Cas9 in quantitative systems biology.

Lipidomics has become a major -omics approach in biomedical research.^10–12^ It can identify thousands of different lipid species, including the structurally diverse membrane lipids. Lipids affect the physicochemical properties of membranes as well as the activity of proteins in contact with them.^13–15^ However, the molecular links between membrane lipid composition and biological processes are still poorly understood. It is therefore critical to accumulate more knowledge about mechanisms regulating membrane lipid composition. The metabolism of membrane lipids is complex due to the interconnection of metabolites from different lipid classes.^13,16^ It has been shown that the change in one lipid can affect the levels of other ones, including those that are metabolically distant, generating a network of co-regulated lipids.^17^ This co-regulation leads to membrane lipid compositions that modulate immune responses upon various stimuli.^17^ We found that sphingolipids (SLs) and ether phosphatidylcholine (ether PC) are co-regulated, which is critical to maintain the integrity of the secretory pathway.^18^ While these studies establish the importance of lipid co-regulation, the mechanisms linking the quantitative change of one lipid to that of another one remains poorly understood. A combination of genetic disruption of lipid-related enzymes and lipidomics should be extremely useful to solve the problem. However, one requisite for such approaches would be that lipid co-regulation is correctly detected in the dataset, which is difficult when using clonal cells as discussed above. Here, we report a strategy that allows us to disrupt genes with an efficiency that is high enough to dispense with clone isolation, which we used to study lipid co-regulation reliably.

## RESULTS AND DISCUSSION

### High mutational activity of CRISPR-Cas9 in a subset of cells

The initial idea of the gene disruption strategy originated from a previous study, in which we established a PCR-based method to detect mutant clones.^19^ In that study, we used CRISPR-Cas9 to mutate three genes (Sgpl1, Sgpp1, and Sgpp2) and analyzed 14 clones to obtain a triple mutant cell line. Only one clone was a full Sgpp2 mutant (no wild type Sgpp2 allele), showing that Sgpp2 sgRNA was ineffective. Nevertheless, the other two targets were fully mutated in this clone. We hypothesized that Sgpp2 disruption occurred in a subset of cells with high mutational activity, in which gene disruption was possible even with the inefficient sgRNA, thus explaining the disruption of the other targets. To test this, we further analyzed 28 clones after transfection with plasmids encoding Cas9 and sgRNAs targeting Sgpl1, Sgpp1, and Sgpp2 (Figure S1A). Transient transfection confirmed that Sgpp2 sgRNA was the least active (Figure S1B). A combination of PCR-based analysis (Figures S1C and S1D) and Sanger sequencing (Figure S1E) detected two mutants of Sgpp2, in which Sgpl1 and Sgpp1 were also mutated (Figures S1E and S1F). Therefore, in all Sgpp2 mutants we obtained, Sgpl1 and Sgpp1 were mutated too. The result strengthened our hypothesis that a subset of cells transfected with plasmids encoding CRISPR-Cas9 components has a high mutational activity.

### Establishment of GENF, a highly efficient CRISPR-Cas9 strategy

In order to take advantage of cells with high mutational activity, we designed a strategy based on HPRT co-targeting,^20^ in which CRISPR-Cas9 is used to mutate Hprt1 together with the intended targets. Hprt1 mutant cells are resistant against the drug 6-thioguanine (6-TG), thus their selection with 6-TG after co-targeting enables the isolation of cells correctly transfected, enriching target-mutated cells.^20^ We hypothesized that the use of conditions where Hprt1 mutation are not efficient, with poor sgRNA design or by transfecting low levels of plasmids encoding Hprt1 sgRNA, 6-TG resistance would be acquired only in cells with high mutational activity (Figure 1A). To test this, we transfected rat McA-RH7777 cells with two plasmids; one (pX330) encoded both Cas9 and the sgRNA against the target gene,^5^ while the other (pHPRTsg) encoded the sgRNA against Hprt1. To alter the efficiency of Hprt1 targeting, we varied Hprt1 sgRNA sequence as well as the ratio between the two plasmids. Consistent with our hypothesis, target (Sphk2) gene mutation rates were highest (99.3%) in 6-TG resistant cells generated under conditions where the efficiency to mutate Hprt1 was the lowest (as reflected by the yield of cells) (Figures 1B and 1C). Thus, via a combination of poor sgRNA design and a low copy number of Hprt1 sgRNA, we achieved nearly complete target mutation after 6-TG selection (Figure 1C). We named this strategy GENF (<u>Ge</u>ne co-targeting with <u>n</u>on-efficient conditions), which mutates genes almost completely without isolating clones. Throughout this study, target mutation rates are calculated from the loss of wild type signals in Sanger sequencing (analyzed by a deconvolution method TIDE^21^) (see validation in Figures S2A-S2C). We further tested GENF in 8 different targets and obtained extremely high target mutation rates (98.6% in the lowest case and 99.9% on average, Figure 1D). Although not being as high as for single target mutation, double or triple gene disruption was also efficient (83.8% in the lowest case and 93.5% on average, Figures 1E and 1F). To test the limit of GENF, we induced 8-plex mutations, for which day-by-day variation became larger (Figure 1G). However, in two out of three experiments high target mutation rates were obtained (71.6% in the lowest case and 92.7% in average). Thus, GENF generates very highly mutated cells under multiplex settings.

**Figure 1.**
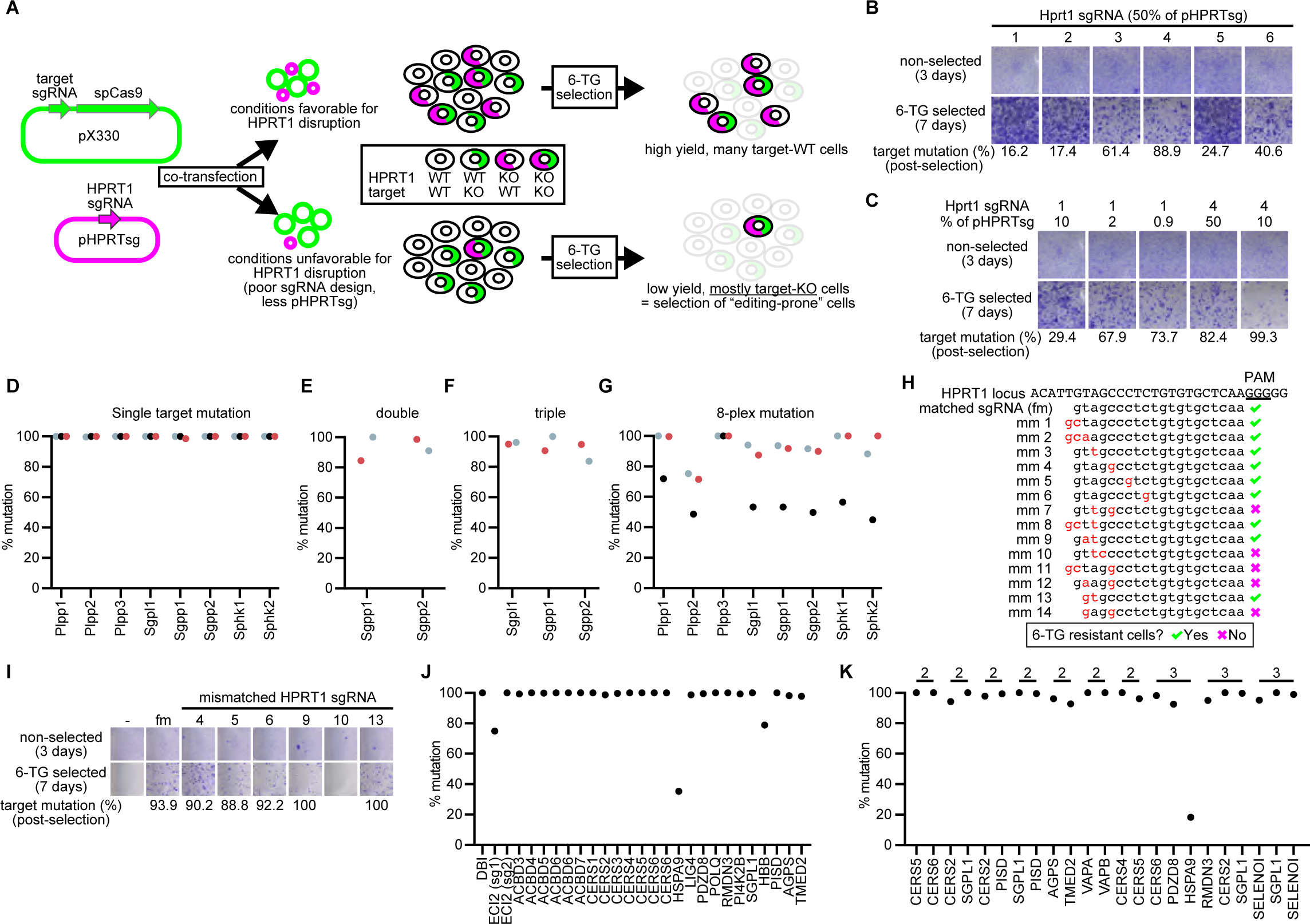
Establishment of GENF. (A) Design of GENF and the plasmids (pX330 and pHPRTsg) used for it. HPRT co-targeting experiments performed under standard conditions^20^ (top) or conditions where Hprt1 disruption is unfavorable (bottom, corresponding to GENF) are illustrated. (B and C) Co-targeting of Sphk2 and Hprt1 in McA-RH7777 cells using different sgRNAs against Hprt1 at various percentage of pHPRTsg plasmid. Cells were stained with Crystal Violet after culture for the indicated time with or without 6-TG selection. The non-selected cells were prepared to measure seeded cell densities. The percentage of Sphk2 mutation (target mutation % post-selection) is shown under the bottom panel. (D-G) Mutation rates after GENF for 8 genes targeted either alone (D), in duplex (E), in triplex (F), or in 8-plex (G). Experiments performed on different days are distinguished by colors. (H) Design of the sgRNA against HPRT1 (sgRNA with a full match, fm) or its 14 variants (mm 1-14) with mismatches (in red) used to establish GENF in HeLa cells. The sgRNAs that enabled the generation of 6-TG resistant cells are labeled with a tick and the others with a cross. PAM: protospacer adjacent motif. (I) Co-targeting of CERS6 and HPRT1 in HeLa cells using representative HPRT1 sgRNAs with (numbers correspond to mm# in (H)) or without (fm) mismatches. 1% (w/w) of pHPRTsg was used. Panels are organized similarly to (B) and (C). (J and K) Mutation rates after GENF for the indicated genes targeted in HeLa cells either alone (J) or in multiplex (K). (K) Targets under the same lines were mutated together. See also Figures S1-S3.

### Mismatched sgRNAs facilitate GENF

When applying the strategy to HeLa cells, we tested multiple HPRT1 sgRNAs but could not obtain low-efficiency ones optimal for GENF. Thus, we investigated whether we could use mismatched sgRNAs to lower HPRT1 mutation efficiencies easily (Figure 1H). Single mismatches in HPRT1 sgRNA did not affect the yield of 6-TG resistant cells and did not increase target mutations in 6-TG resistant cells (Figure 1I). On the other hand, sgRNAs with two mismatches enabled the generation of 6-TG resistant cells only when the mismatches were in the distal-most positions from the NGG proximal adjacent motif (Figures 1H and 1I). Such sgRNAs enabled complete post-selection target gene (CERS6) mutations (Figure 1I). Extremely high gene disruption was achieved with this protocol in multiple targets, including in multiplex experiments (Figures 1J and 1K). One of the few targets we could not mutate efficiently with GENF was HSPA9, which is an essential gene according to DepMap database^22^ (Figure S3A). We note that another essential gene, TMED2, could be mutated with GENF, but with indels strongly enriched for in-frame mutations (Figures S3A and S3B). The other two targets with low mutation rates were ECI2 and HBB (Figures 1J and 1K). Transient transfection experiments revealed that the sgRNAs for these targets were poorly active (Figure S3C). Therefore, GENF enables highly efficient gene disruption unless the sgRNA is poorly active or when the target is essential.

### Additional drug-gene pair for GENF

One limitation of GENF was that it cannot be repeated sequentially on the modified cells. To overcome this, we looked for additional drug-gene pairs to be used for GENF from published genome-wide CRISPR screenings that study drug resistance. A study reported that POR (encoding cytochrome P450 reductase) mutation confers resistance against paraquat,^23^ thus we tested this drug-gene pair. POR targeting led to the formation of small colonies after paraquat treatment (Figures 2A and 2B). Performing POR co-targeting using an sgRNA with two mismatches enabled a reproducible near-complete depletion of a target gene (CERS6), thus establishing an additional drug-gene pair for GENF (Figures 2A-2C). The small size of colonies is consistent with a partial toxicity of paraquat in POR mutant cells,^23^ thus we recommend the use of HPRT1 primarily for GENF. Having an additional drug-gene pair for GENF, we could repeat gene disruption two times. Using this, we could overcome the slightly lower efficiency of highly multiplexed gene disruption with GENF (Figures 2D-2F). Using HPRT1 as the co-targeted gene, we generated with GENF a cell line having mutations in all six CERS genes (designated ΔCERS) with high but imperfect efficiencies (88.5% for the lowest case and 93.4% on average, Figure 2F). We targeted again the six CERS genes in ΔCERS cells with GENF, using POR as the co-targeted gene and paraquat for selection (Figure 2E). We obtained a cell line (designated Δ/ΔCERS) having near-complete mutations in all CERS genes (95.9% for the lowest case and 99.2% in average, Figures 2D-2F). Although CERS genes are critical for the synthesis of complex sphingolipids, Δ/ΔCERS cells could be maintained in cultured similarly to control cells, probably due to the supply of lipids from the serum (as discussed later).

**Figure 2.**
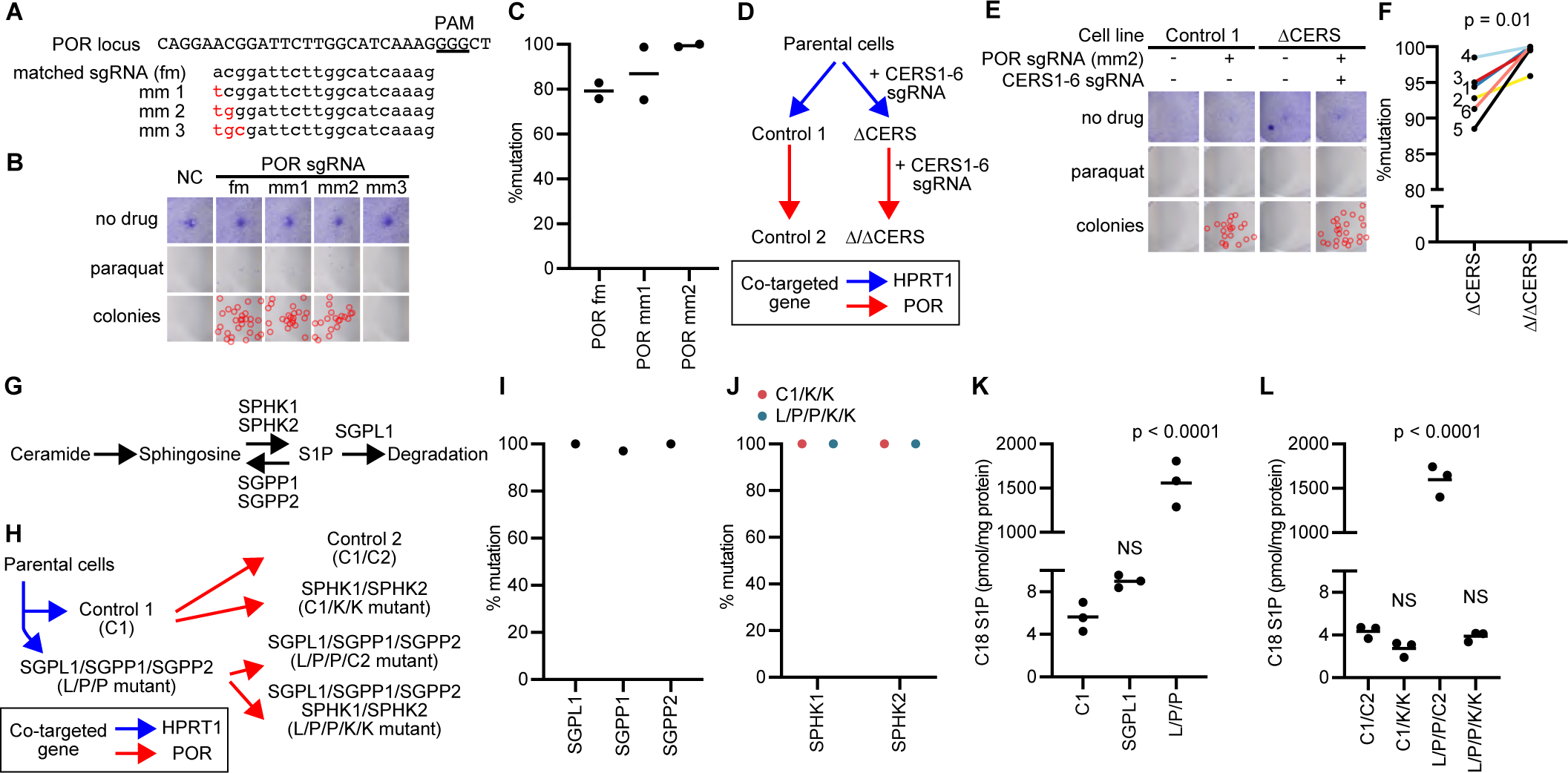
Establishment of GENF with another gene-drug pair. (A) Sequence of the targeted POR locus and the sgRNAs targeting it with (mm) or without (fm) mismatches at PAM-distal positions. (B) Cells were stained with Crystal Violet after co-targeting POR and CERS6, with or without treatment with paraquat. The colonies of drug-resistant cells are highlighted with circles in additional panels, due to their smaller size than those obtained after HPRT co-targeting. (C) Mutation rates of CERS6 after co-targeting with the indicated sgRNA and selection with paraquat in two independent experiments. (D) Nomenclature of cell lines obtained during two-step GENF, with the different co-targeted genes color coded. (E) Survival of cells after the second round of two-step GENF (corresponding to the red arrows in (D)), which was analyzed similarly to (B). Note the absence of paraquat-resistant cells when POR was not co-targeted. (F) Mutation rates of CERS genes (see numbers) all targeted together, in the first step (ΔCERS) or the second step (Δ/ΔCERS) of GENF. See the increase in mutation rates for all targets. p value: paired t-test. (G) The metabolic pathway that was manipulated in another experiment of two-step GENF. Sphingosine is generated through degradation of ceramide and is phosphorylated into sphingosine 1-phosphate (S1P) by sphingosine kinases (SPHK1, SPHK2). S1P is dephosphorylated by S1P phosphatases (SGPP1, SGPP2) or degraded by S1P lyase (SGPL1). (H) Combination of genes mutated during two-step GENF, together with the nomenclature of the mutants. (I) Mutation rates in L/P/P mutants generated in the first step illustrated in (H). (J) Mutation rates of genes targeted in the second step illustrated in (H). (K and L) S1P levels in mutants generated in the first (K) and second (L) steps. p values: one-way ANOVA followed by Dunnett’s multiple comparison test to compare with controls C1 (K) or C1/C2 (L).

In another experiment, we manipulated metabolite levels with GENF and reverted the phenotypes by adding mutations through another round of GENF. We generated a cell line (designated L/P/P) lacking SGPL1, SGPP1, and SGPP2, which are enzymes that reduce the amount of sphingosine 1-phosphate (S1P) through dephosphorylation or degradation (Figure 2G).^24^ From this cell line and a control cell line (with only HPRT1 mutated, designated C1), we generated cell lines with mutations in the two kinases (SPHK1 and SPHK2) that produce S1P (Figures 2G and 2H). L/P/P cells were generated with high mutation efficiency in the first round of GENF (97.1% for the lowest case and 99.0% on average, Figure 2I). In the second round, both SPHKs were completely mutated in L/P/P cells, generating L/P/P/K/K cells, or in the control cell line C1, generating C1/K/K cells (Figures 2H and 2J). Control cells of this second step, in which only POR was mutated, were designated C1/C2 and L/P/P/C2 cells (Figure 2H). S1P levels in L/P/P cells were drastically increased (>270 fold), while SGPL1 mutation alone had only a marginal effect (Figure 2K). Importantly, L/P/P/K/K cells had comparable S1P levels to control cells or C1/K/K cells, demonstrating that phenotypes of mutant cells can be reverted through repeated GENF (Figure 2L). We therefore made it possible to repeat GENF to further enhance multiplex targeting efficiencies, or to alter phenotypes of established mutants.

### Characterization of mutations obtained with GENF

CRISPR-Cas9 is known to have some limitations, such as the variability in on-target effects and the presence of off-target effects.^25,26^ GENF was designed specifically to address the issues related to clonal variability, thus it did not solve these other limitations. Nevertheless, it was important to characterize them. We tested the on-target mutation patterns using four mutant McA-RH7777 cell lines lacking either Agpat1 or Agpat2, generated with two different sgRNAs each, each generated in triplicate. While different sgRNAs led to diverse indel patterns, different cell lines generated with the same sgRNA had very similar patterns (Figures 3A and 3B). Indel patterns were also reproducible when comparing cell lines generated on different days (Figure S4A). This showed that GENF generates reproducible on-target effects from the same sgRNA. Our analysis was not suitable for detecting kb-sized large-scale indels as well as transpositions or inversions, all having been reported with CRISPR-Cas9 experiments.^25^ However, extrapolating from the reproducible patterns seen in our experiments, it is likely that these adverse on-target effects are also happening to a similar extent when using the same sgRNA. We also note that such adverse mutations should be diluted in the polyclonal population when using GENF, while they should be present stochastically when using clones.

**Figure 3.**
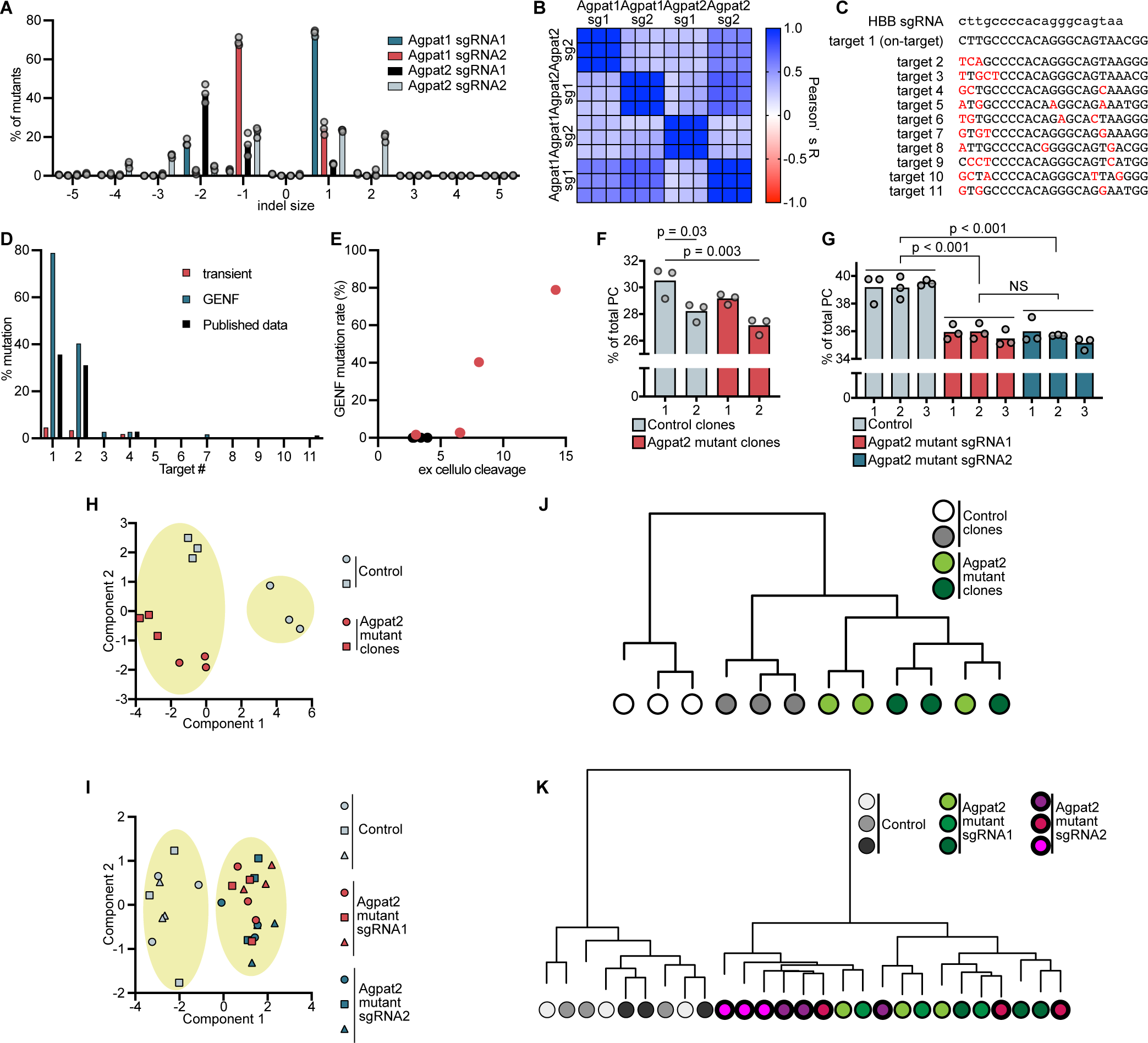
Characterization of mutants generated by GENF. (A and B) Indel patterns seen in 12 mutant cell lines generated with four different sgRNAs, and similarity between them assessed by Pearson’s correlation coefficients. (C-E) Analysis of off-target mutations in HBB mutant cells. (C) Off-target candidates of HBB sgRNA reported from *ex cellulo* cleavage experiments, with mismatches highlighted in red.^27^ (D) On- and off-target mutations in HBB mutant cells generated with GENF, as compared to those obtained in a published study.^27^ Mutations obtained by transient transfection are also shown to demonstrate the enrichment obtained by GENF. (E) On- and off-target mutation rates in HBB mutants compared with published *ex cellulo* cleavage results.^27^ Non-zero values are plotted in red. Note that mutation rates are well predicted from *ex cellulo* cleavage. (F-K) Comparison of lipid analysis datasets obtained from clones and polyclonal cells generated with GENF. (F and G) Levels of the most abundant phosphatidylcholine (PC) species, PC 34:1, in different control and Agpat2 mutant clones (F) or in different polyclonal cell lines generated with GENF (G). p values: one-way ANOVA followed by Tukey’s multiple comparisons test comparing all pairs. (H and I) Principal component analysis of PC acyl-chain composition datasets obtained from clonal cell lines (H) or polyclonal cell lines generated with GENF (I). Encircled are the results of K-means clustering to split datasets into two groups. (J and K) Hierarchical clustering to assess the similarity of the same datasets from clonal cell lines (J) or polyclonal cell lines generated with GENF (K). See also Figure S4.

We next analyzed off-target effects using the HBB mutant cell line generated with GENF. This cell line was generated with an sgRNA having a well characterized off-target candidate list (Figure 3C), which was obtained from cleavage patterns of genomic DNA preparations from cells (*ex cellulo* cleavage).^27^ We found that the off-target effects of GENF (Figure 3D), remained predictable from *ex cellulo* cleavage (Figure 3E). We also analyzed off-target cleavage in CERS4 mutant cells, which was generated using an sgRNA with the lowest predicted specificity score^5^ among those used in this study. Off-target mutations were present only in targets with three mismatches and not in those with four mismatches (Figure S4C). In addition, the off-target mutation rates were lower than the on-target 100% mutation (Figure S4C). Therefore, GENF has off-target effects similar to other CRISPR-Cas9 protocols, but the off-targets remain predictable, and might be prevented if using sgRNAs that are specific enough. We note that the Cas9 protein used in this study was from the first generation, and “high fidelity” versions^28^ might improve the specificity of GENF. We did not test this possibility in this study, because the use of mismatched HPRT1 sgRNAs would be impossible.

### GENF reveals lipid changes associated with the loss of a disease-related enzyme

Next, we investigated if the use of polyclonal mutant cells generated with GENF improves phenotype analyses and answers biological questions by eliminating clonal variabilities. For this, we analyzed lipid regulation by Agpat2, a lipid-related enzyme mutated in patients with congenital generalized lipodystrophy type 1^29^ (CGL1). The lipid changes that cause CGL1 remain obscure, as different studies report inconsistent lipid alterations associated with Agpat2 loss.^30–32^ We previously found that AGPATs regulate phosphatidylcholine (PC, the most abundant glycerophospholipid) acyl-chains and membrane physical properties.^33,34^ Therefore, we analyzed PC acyl-chains in Agpat2 mutant McA-RH7777 cells generated either by clone isolation (generated in parallel to a previous study^35^) or by GENF. Consistent with the literature describing phenotypic variabilities between clones,^6–9^ we found a variability in PC acyl-chains between clones (Figures 3F and S4D). In contrast, differences between polyclonal cell lines were smaller, and we could find a distinct PC acyl-chain profiles between control and Agpat2 mutant cells (Figures 3G and S4E). We performed multivariate analyses to compare the performance of both approaches (clone isolation versus GENF) for detecting lipid changes induced by gene disruption. Principal component analysis (PCA) could not discriminate between control and Agpat2 mutant clones with the first dimension, while mutants generated with GENF were clearly discriminated (Figures 3H and 3I). When splitting the datasets into two groups with K-means clustering, one control clone was clustered together with the two Agpat2 mutant clones (Figure 3H), while GENF-generated control and Agpat2 mutant cells were discriminated (Figure 3I). Finally, when we assessed data similarity using hierarchical clustering, clones could not be correctly separated based on Agpat2 genotypes, while cell lines generated by GENF were clearly distinguished (Figures 3J and 3K). Consequently, we were able to discover lipids regulated by Agpat2 only when using polyclonal mutants generated by GENF, which shows that polyclonality is critical to obtain reliable quantitative datasets.

Since standard culture conditions (medium supplemented with 10% fetal calf serum) are poor in polyunsaturated fatty acid (PUFA), making cell lines PUFA-poor,^36,37^ we investigated PC acyl-chains of cells supplemented with three PUFAs: linoleic acid, arachidonic acid, and docosahexaenoic acid. In such conditions, Agpat2 mutation mainly decreased PC 34:2 in the mutants (Figure 4A). MS/MS fragmentation revealed that this decrease is attributable to reductions in linoleic acid (PUFA with 18 carbons and 2 double bonds) (Figures S5A-S5C). Gas chromatography analysis confirmed that linoleic acid levels are decreased in glycerophospholipids of Agpat2-mutant cells (Figure 4B). Since Agpat2 encodes an enzyme with lysophosphatidic acid acyltransferase (LPAAT) activity, we analyzed how Agpat2 disruption affects this activity and its substrate preference. Agpat2 mutants had decreased LPAAT activity (Figure 4C) as well as the preference for linoleic acid incorporation (Figure 4D and Figures S5D-S5G). Our previous study suggested that LPAAT activity regulates linoleic acid levels in PC^33^ and our results confirmed this observation while identifying Agpat2 as the responsible enzyme. We compared the selectivity of LPAAT activity and gene expression profiles of Agpat enzymes in various tissues and found that tissues with high Agpat2 expression have an LPAAT activity with higher preference for linoleic acid (Figure S5H). In addition, cells overexpressing Agpat2 had increased PC 34:2 levels (Figure 4E). All these results establish Agpat2 as an enzyme regulating linoleic acid levels in PC. Thus, GENF revealed a potential link between membrane lipid acyl-chains and a genetic disease with impaired adipogenesis, which will be investigated in the future.

**Figure 4.**
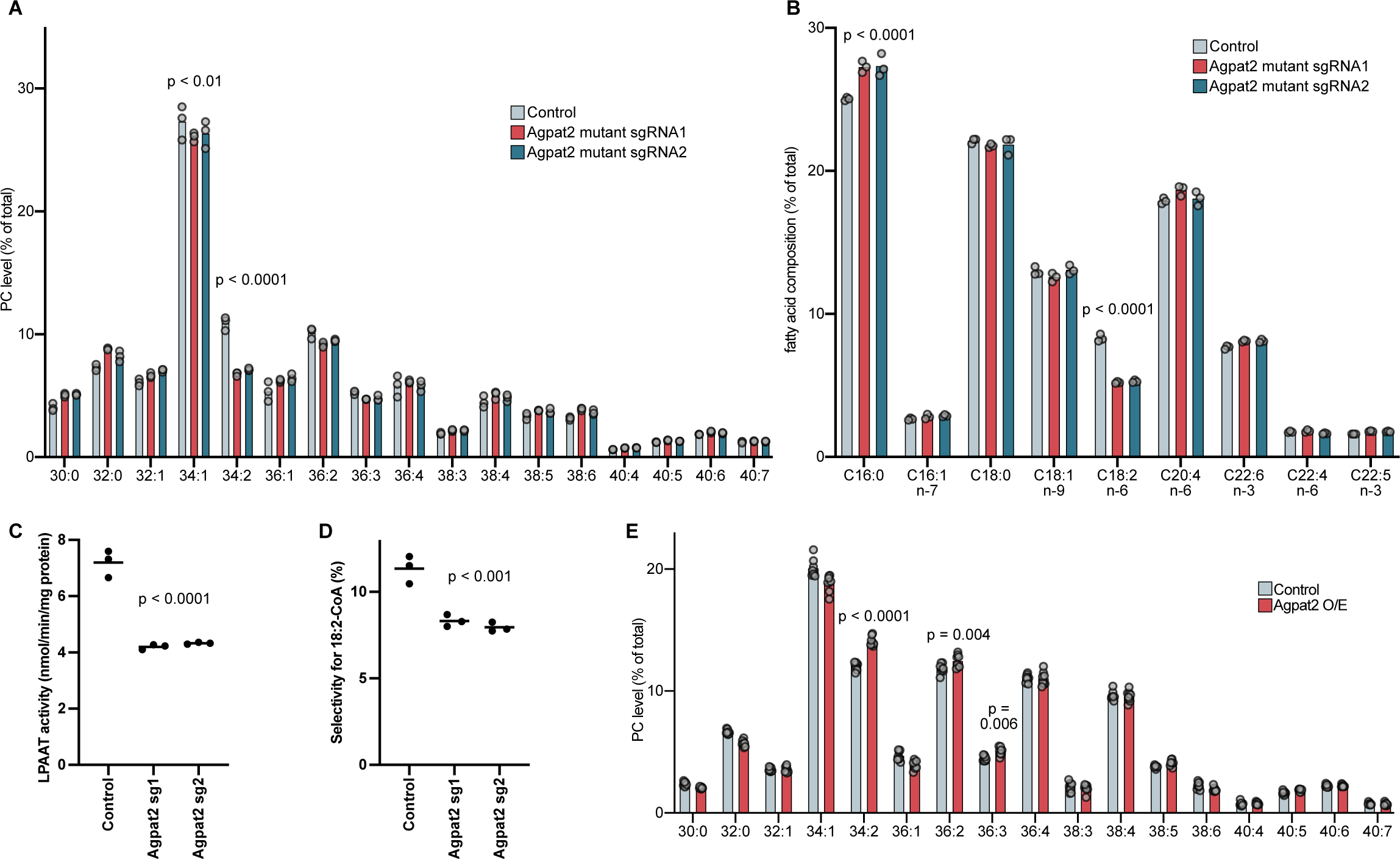
Agpat2 regulates linoleic acid levels in PC. (A) PC acyl-chain composition in control or Agpat2 mutant cells supplemented with linoleic, arachidonic, and docosahexaenoic acid (PUFA supplementation). p values: two-way ANOVA followed by Dunnett’s multiple comparisons test to compare with Control. Only reductions seen in both Agpat2 mutants are illustrated for simplicity. (B) Gas chromatography analysis of fatty acid methyl esters obtained from phospholipid fractions of PUFA-supplemented control or Agpat2 mutant cells. p values: two-way ANOVA followed by Dunnett’s multiple comparisons test to compare with Control. Common changes are illustrated. (C and D) Lysophosphatidic acid acyltransferase (LPAAT) activity, and its selectivity for linoleoyl-CoA in membrane fractions obtained from control or Agpat2 mutant cells. p values: one-way ANOVA followed by Dunnett’s multiple comparisons test to compare with Control. (E) PC acyl-chain composition in PUFA-supplemented control or Agpat2-overexpressing (O/E) cells. p values: two-way ANOVA followed by Sidak’s multiple comparisons test. Only increases are illustrated for simplicity. See also Figure S5.

### Manipulation of sphingolipid acyl-chains with GENF

Using GENF, we generated a series of mutant cells lacking individual ceramide synthases (CERS1-CERS6), which incorporate N-acyl-chains in sphingolipids (SLs)^38^ (Figure 1J). Published RNA-seq data of the HeLa clone that we use^39^ revealed that only CERS2 and CERS5 are expressed in the parental cell line (Figure S6A). Therefore, mutants of the four non-expressed CERSs should not have any phenotypes unless GENF generated artifacts. We performed lipidomics analyses of the mutant cell lines to investigate changes in SLs and glycerophospholipids. CERS2 mutants had drastic decreases in C24 ceramide (C24 Cer) and compensatory increases in C16 ceramide (Figures 5A-5C). The same pattern was seen in the complex SLs sphingomyelin (SM) and hexosylceramide (HexCer) (Figures 5D-5F and S6B-S6D). On the other hand, CERS5 mutant had decreased C16 ceramide and C16 SM levels, while other mutants had no changes in SL N-acyl-chains (Figures 5A, 5B, 5D, 5E, S6B, and S6C). The results show that CERS2 and CERS5 compete for the same substrate, and that the loss of CERS2 leads to an increased flux of sphingosine usage by CERS5 (Figures 5G and 5H). Indeed, the increase of C16 SLs was minimal in Δ/ΔCERS cells mutated for all CERSs, despite drastic reductions in C24 SLs (Figures 5I, 5J, and S6E). We note a considerable amount of C16 SLs detectable in Δ/ΔCERS cells, which is probably due to the supply of SLs from the serum added to the culture medium, as has been documented^40^. The reverse reaction of ceramidases has also been reported to generate ceramides^41^. Despite the drastic reductions in SLs seen in Δ/ΔCERS cells (Figure 5K), the cells could be maintained in culture similarly to control cells with similar passage frequencies, showing the ability of cell lines to survive with SLs provided from the culture medium.

**Figure 5.**
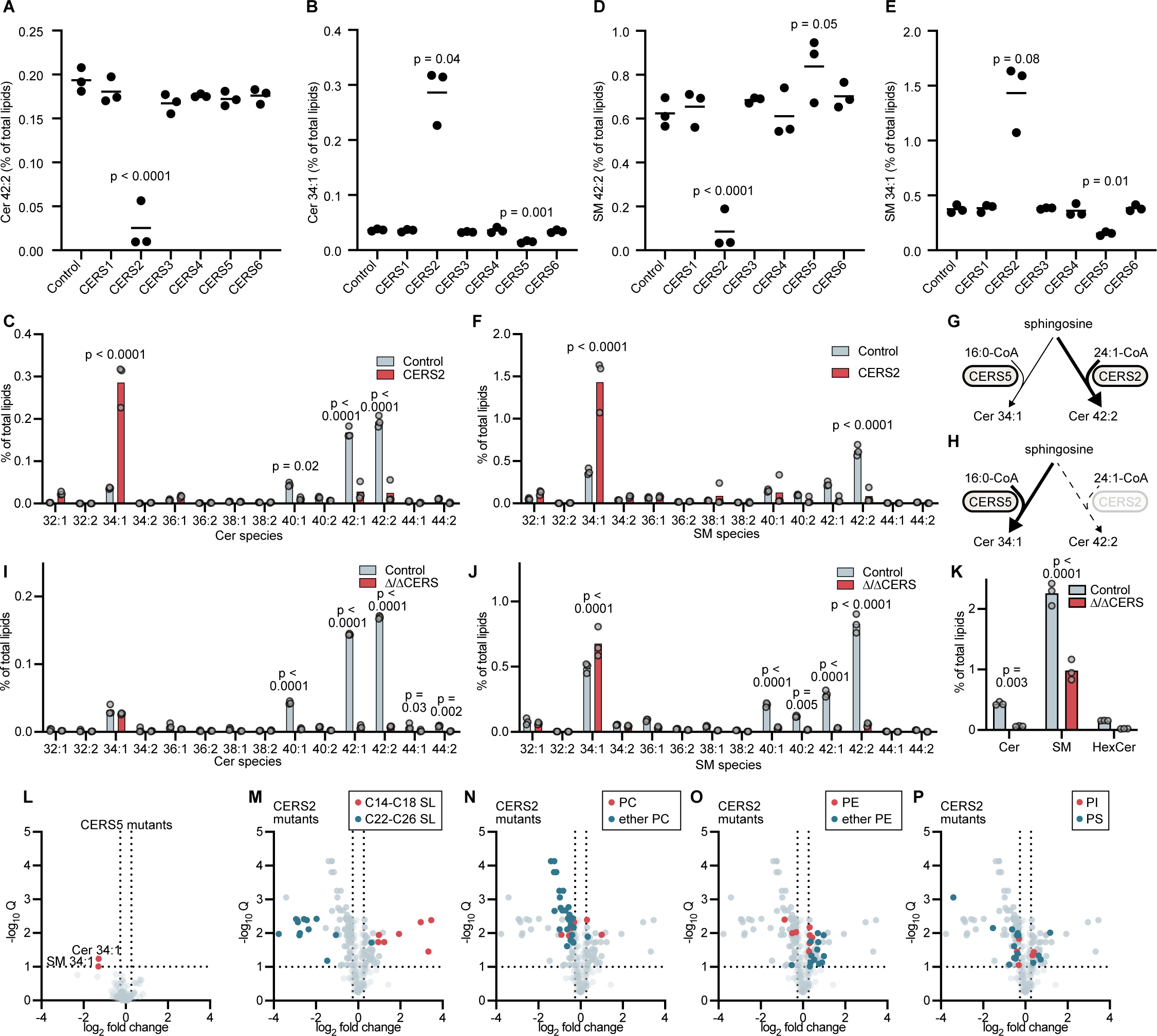
Lipidome changes induced by mutations in CERSs. (A and B) Levels of the indicated ceramide (Cer) species in various CERS mutants. (C) Changes in Cer species in CERS2 mutants. (D and E) Changes in the indicated sphingomyelin (SM) species in various CERS mutants. (F) Changes in SM species in CERS2 mutants. (G and H) The competition between CERS2 and CERS5 explains the changes in SL profiles seen in CERS2 mutants. (I and J) Changes in Cer (I) and SM (J) species in Δ/ΔCERS mutants. (K) Changes in total levels of the indicated sphingolipid species in Δ/ΔCERS mutants. p values: (A and D) one-way ANOVA followed by Dunnett’s multiple comparisons test to compare with Control, (B and E) Brown-Forsythe and Welch ANOVA tests followed by Dunnett’s T3 multiple comparisons test to compare with Control, (C, F, and I-K) two-way ANOVA followed by Sidak’s multiple comparisons test. (L-P) Volcano plot illustrating changes in lipid levels and their statistical significances in the indicated CERS mutants. Lipids with changes above 1.2-fold and q values (FDR-corrected multiple t-test) below 0.1 were considered as hits. (M-P) For CERS2 mutants, different lipid classes are highlighted separately. See also Figure S6.

We further analyzed lipidomics data of single CERS mutants. Consistently with the lack of their expression in parental cells, CERS1, CERS3, CERS4, and CERS6 mutants had no significant changes in all lipids analyzed (Figures S6F-S6I). It is unlikely that clonal mutants, with their high variances, could lead to similar unchanged datasets comprising >230 lipids. CERS5 mutants had only two significantly changed lipids, both being C16 SLs (Figure 5L). Unexpectedly, CERS2 mutants had a drastic rearrangement of the whole lipidome (Figures 5M-5P). In addition to the changes in SLs (Figure 5M), we found that ether PC, with ether or vinyl ether bonds at the *sn-*1 position, is decreased in CERS2 mutants (Figure 5N). Multiple phosphatidylethanolamine (PE) species were increased, but the effect was stronger with PE having an ether or vinyl-ether bond at the *sn-*1 position (ether PE) (Figure 5O). Some decreases in phosphatidylserine (PS) species were also seen (Figure 5P). This revealed a whole lipid co-regulatory network associated with a decrease in C24 SL production. Our results show the power of GENF to generate mutants and manipulate metabolite levels, and to detect co-regulatory networks associated with the manipulations. It is important to note that the data reliability of GENF is especially critical to analyze co-regulatory networks. With clonal cells, variabilities in datasets would make it difficult to discriminate whether the detected changes reflect a co-regulation with the metabolites that we manipulated or they are accidental changes due to clonal variations.

### Dissection of co-regulatory networks with multiplex GENF

The rapidity and efficient multiplexing of GENF accelerates the process of hypothesis refinement, as it allows us to remake new mutants rapidly after experimental data suggest new hypotheses. Here, we used GENF to analyze the mechanisms of ether lipid changes in CERS2 mutants, due to the importance of the co-regulation between SLs and ether PC to maintain the integrity of the secretory pathway.^18^ CERS2 mutants had decreased ether PC, with strong increases in ether PE and small increases in PE (hereafter, PE denotes non-ether PE) (Figures 6A-6C). The sum of ether PC and ether PE was unchanged (Figures 6D), suggesting that the balance of head group incorporation into the common precursor, ether diacylglycerol (ether DAG) was affected in CERS2 mutants (Figure 6E). At the level of individual species, most ether PC species (saturated, monounsaturated, and polyunsaturated) were decreased in CERS2 mutants, while only polyunsaturated ether PE species increased (Figures 6F and 6G).

**Figure 6.**
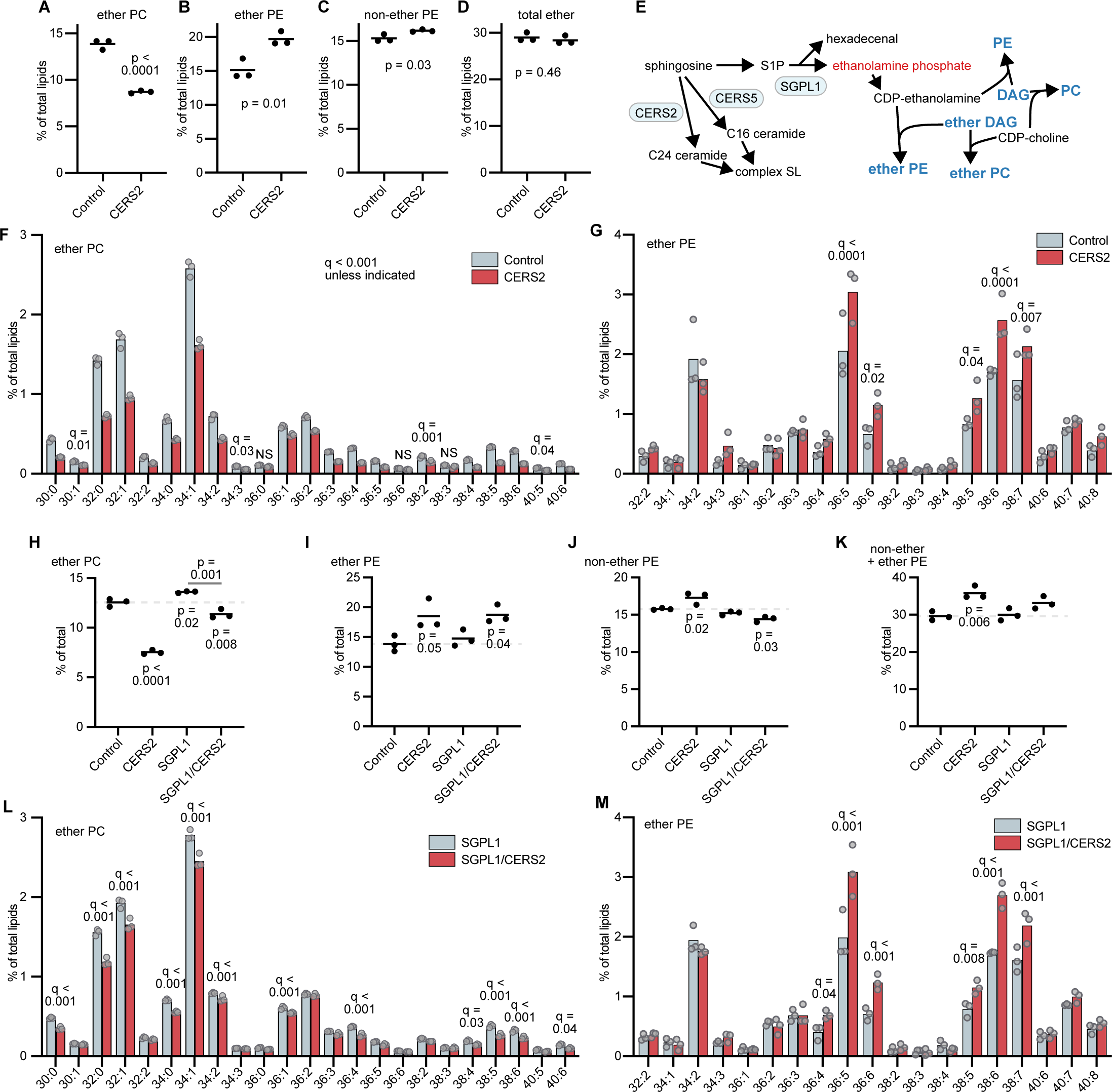
SL and ether lipid co-regulation through SL degradation. (A-D) Changes in the indicated lipid classes in CERS2 mutants. p values: unpaired t-test. (E) Metabolic pathways linking SL degradation and synthesis of PE or ether PE. (F and G) Changes in ether PC and ether PE species in CERS2 mutants. q values: one-way ANOVA followed by 5% FDR-corrected multiple t-test. (H-K) Changes in the indicated lipid classes in the indicated mutants. p values: one-way ANOVA followed by followed by Sidak’s multiple comparisons test. Values below the bars are comparisons with Control, and those above are comparisons between indicated pairs. (L and M) Changes in ether PC and ether PE species in CERS2 mutants in an SGPL1 mutant background. q values: two-way ANOVA followed by 5% FDR-corrected multiple t-test.

One degradation product of SLs, ethanolamine phosphate, is used for PE and ether PE synthesis after conversion into CDP-ethanolamine^24^ (Figure 6E). We thus hypothesized that CERS2 mutation increased the flux of sphingosine toward degradation, upregulating ethanolamine phosphate production for PE and ether PE synthesis, at the expense of ether PC. We generated a series of cells with various combinations of mutations in CERS2 or SGPL1, the enzyme responsible for SL degradation^24^ (Figures 1J, 1K, and 6E). In contrast to the large decrease seen in CERS2 single mutants, ether PC levels were close to control cells in SGPL1/CERS2 mutants (Figure 6H). As compared to control cells, SGPL1/CERS2 mutants had higher and lower ether PE and PE, respectively, making the sum of PE and ether PE almost similar to controls (Figures 6I-K). Thus, SGPL1-dependent ethanolamine phosphate production contributes to the increases in total PE (PE plus ether PE) and the decreases in ether PC under CERS2 deficiency.

Nevertheless, an additional layer of CERS2-dependent ether lipid regulation was suggested from a comparison between SGPL1 and SGPL1/CERS2 mutants. SGPL1/CERS2 mutants had lower ether PC and a tendency for higher ether PE than SGPL1 mutants (Figures 6H and 6I). Individual ether PC species decreased by CERS2 mutation to a lesser extent in an SGPL1 mutant background than in a wild type background (Figures 6F and 6L). Most saturated and only a few polyunsaturated ether PC species decreased by CERS2 mutation in an SGPL1 mutant background (Figure 6L). The ether PE species that increased by CERS2 mutation were polyunsaturated, irrespective of the SGPL1 genotype (Figures 6G and 6M). Among the two enzymes that synthesize PE and ether PE (SELENOI and CEPT1, Figure 7A), SELENOI contributes more to ether PE synthesis with preferences for polyunsaturated substrates.^42,43^ To test whether SELENOI contributed to the increases of ether PE in SGPL1/CERS2 mutants, we mutated SELENOI and CERS2 alone or in combination, all in an SGPL1 mutant background (Figure 1K). SELENOI mutations increased ether PC and decreased ether PE (Figures 7B and 7C). CERS2 mutation decreased ether PC both in SGPL1 mutant and SGPL1/SELENOI mutant background (Figure 7B), while it increased ether PE in SGPL1 mutant but not in SGPL1/SELENOI mutant background (Figure 7C). This showed that SGPL1-independent increases in ether PE upon CERS2 mutation are SELENOI-dependent. At the molecular species level, CERS2 mutation did not decrease polyunsaturated ether PC anymore in an SGPL1/SELENOI mutant background (Figure 7D). Nevertheless, CERS2 mutation still decreased saturated and monounsaturated ether PC and increased polyunsaturated ether PE at the expense of saturated and monounsaturated ether PE in SGPL1/SELENOI mutant background (Figures 7D and 7E). This suggests that CERS2 mutation causes additional SELENOI/SGPL1-independent decreases in saturated and monounsaturated ether PC and ether PE, which we will investigate in the future.

**Figure 7.**
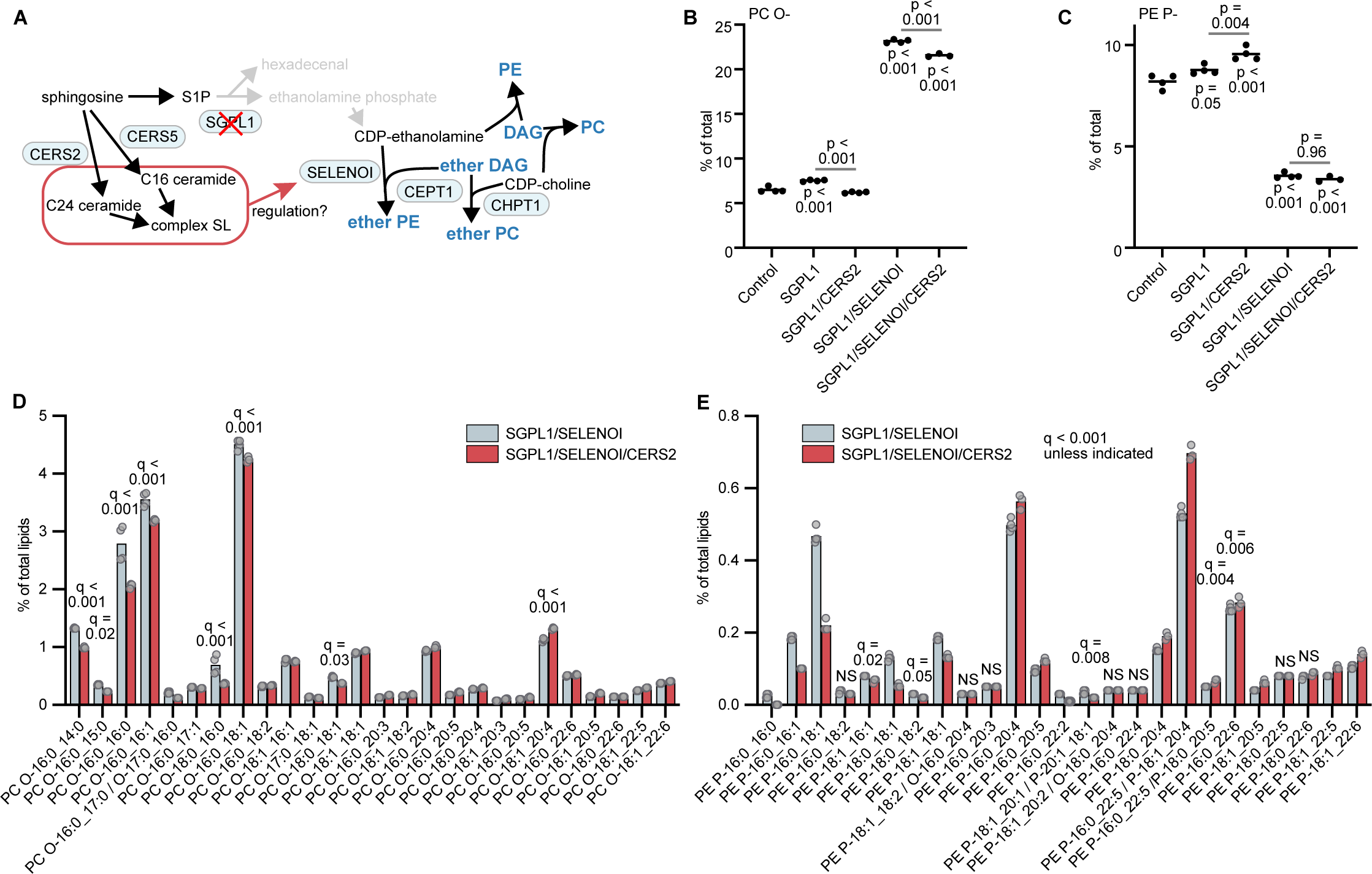
SELENOI-dependent co-regulation between SL and ether lipids. (A) Hypothesis about ether PE upregulation by CERS2 mutation in the absence of SGPL1. (B and C) Ether PC and ether PE levels in the indicated mutants. p values: one-way ANOVA followed by followed by Sidak’s multiple comparisons test. Values below the bars are comparisons with Control, and those above are comparisons between indicated pairs. (D and E) Changes in ether PC and ether PE species in CERS2 mutants in an SGPL1/SELENOI mutant background. q values: two-way ANOVA followed by 5% FDR-corrected multiple t-test.

Thus, the rapidity of GENF allowed us to repeat multiple rounds of hypothesis testing, which revealed three layers of ether lipid changes that occur in CERS2 mutants. SGPL1-dependent SL degradation increases both ether PE and PE, and decreases ether PC. SELENOI-dependent ether PE synthesis is promoted, thereby decreasing polyunsaturated ether PC due to the consumption of polyunsaturated ether DAG. Moreover, SGPL1/SELENOI-independent changes occur in CERS2 mutants, resulting in the reduction of saturated and monounsaturated ether PC and ether PE.

### Conclusions, limitations, and future directions

We developed GENF as a strategy for cloning-free gene mutation. While GENF is based on the selection of cells that can mutate their targets even with a non-efficient sgRNA, the precise mechanism of how it works remains unknown. One mechanism could be that we are selecting cells that received more plasmids and expressed higher amounts of Cas9 and sgRNAs. However, in a study using cell sorting to isolate transfectants with high Cas9 levels, gene disruption was efficient but did not reach 100%.^44^ In contrast, when using GENF non-disrupted alleles were barely detected. Thus, it is likely that GENF is not solely based on the selection of high expressors or CRISPR-Cas9 components, but rather involves other factors such as the epigenetic state of the genome. It should be noted that some methods achieve very good mutation rates with CRISPR-Cas9 without clone isolation.^45,46^ One benefit of GENF is that, once optimized, it is a simple method involving only transfection, drug selection, and occasional passages. This simplicity makes it very easy to generate multiple mutant cell lines in parallel in a time- and cost-efficient way.

The biggest advantage of GENF (and other polyclonal strategies) is the avoidance of artifacts related to clonal variances. The discovery of PC acyl-chains regulated by Agpat2 provides a good example of a finding that would be difficult to make with clonal cell lines. The use of polyclonal cell lines is recommendable in all experiments where control and mutant cells are compared. We note that polyclonal mutant cells have already been used in previous studies, in which mutants were generated by lentiviral delivery of CRISPR-Cas9 components and drug selection of transduced cells.^47^ While such approaches could be a good alternative to GENF, some studies revealed that the transduced cell populations are not always fully mutated.^48^ This incompleteness could be problematic especially when multiplex gene disruption is performed to revert phenotypes, as has been done in our analysis of the co-regulation of ether lipids and SLs. In addition, GENF is a virus-free approach, enabling its use in most laboratories.

Through a combination of highly efficient gene disruption enabled by GENF and lipidomics analyses, we obtained reliable datasets to propose novel concepts in lipid biology. Based on the continuity from our previous research,^33,35^ we mainly focused on the roles of membrane lipid acyl-chains and revealed a potential role of reduced linoleic acid-containing glycerophospholipids in the genetic disease congenital generalized lipodystrophy type 1. We also found that the N-acyl-chains of SLs regulate the metabolism of ether lipids in a complex manner, with at least three different mechanisms. The dissection of these multiple layers of SL-ether lipid co-regulation was possible thanks to the efficient multiplex mutations and reliable lipidomics data that are obtained with GENF.

While we used GENF to study lipids, the method is not limited to this purpose. It is likely that the majority of quantitative traits of cells are affected by clonal variances, and thus the accuracy of studies using mutant clones generated with CRISPR-Cas9 might be affected, if not erroneous. GENF was specifically developed to avoid artifacts caused by clonal variances, which is a severe limitation of CRISPR-Cas9 if combined with clone isolation. Indeed, while the danger of using clones has already been discussed,^6–9^ a large fraction of studies still uses isolated clones. GENF should be especially useful for -omics studies that analyze many variables, such as transcriptomics or metabolomics. On the other hand, we note that GENF does not solve all the limitations of CRISPR-Cas9, such as off-target effects^26^, deleterious on-target mutations^25^, in-frame mutations, or the limitation in target sites due to restrictions of protospacer adjacent motifs^49^. However, researchers have paid more attention to these limitations, and strategies are being developed to solve them, such as the development of improved Cas9 variants^28^. A combined use of GENF with Cas9 variants will be an interesting approach for the future.

## ACKNOWLEDGEMENTS

We are grateful to all members of the Riezman laboratory, Shimizu laboratory, Antonny laboratory, and Harayama laboratory for valuable comments and continuous support. We thank Hideo Shindou (National Center for Global Health and Medicine) for providing research space and advices, Noemi Jiménez-Rojo (University of Geneva) for constructing some of the pX330-based plasmids and sharing preliminary observations, Isabelle Riezman (University of Geneva) for all technical assistance, Keisuke Yanagida (National Center for Global Health and Medicine) and Daisuke Hishikawa (Nippon medical school) for critical reading of the manuscript, Hiroshi Tsugawa (Tokyo University of Agriculture and Technology) for support on lipid analysis by MS-DIAL5, and Nathalie Leroudier for Sanger sequencing analysis. T.H. was supported by the Japan Society for the Promotion of Science (JSPS) Postdoctoral Fellowships for Research Abroad, the French Government (National Research Agency, ANR) through the “Investments for the Future” programs LABEX SIGNALIFE ANR-11-LABX-0028 and IDEX UCAJedi ANR-15-IDEX-01, the ATIP-Avenir program (CNRS/Inserm), and by the Global Innovation Research funds of Tokyo University of Agriculture and Technology. T.H.-Y. was supported by MEXT/JSPS KAKENHI grant 16K21651. T.S. was supported by the AMED Gapfree Program and Takeda Science Foundation 15668360. H.R. was supported by the Swiss National Science Foundation and the NCCR Chemical Biology (grants 184949 and 185898).

## AUTHOR CONTRIBUTIONS

T.H. designed the project and performed data analysis. T.H., T.H.-Y., A.A.-R., K.R., and R.M. generated the mutant cells. T.H., T.H.-Y., L.F., F.H., and D.D. did lipid analysis. T.H., T.S., and H.R. acquired funding and supervised the project. R.M. and A.A.-R. obtained preliminary data for the project. All authors contributed to the writing of the manuscript.

## METHODS

### Cell culture

McA-RH7777 cells were obtained from ATCC (CRL-1601) and HeLa cells (MZ clone) were kindly provided by Jean Gruenberg (University of Geneva). Both cell lines were cultured in Dulbecco’s Modified Eagle Medium (DMEM high glucose, with GlutaMAX and pyruvate), supplemented with 10% fetal bovine serum and contamination preventives (100 units/mL penicillin + 100 µg/mL streptomycin (Thermo Fisher Scientific) or 1x ZellShield (CliniSciences). McA-RH7777 were cultured on collagen-coated dishes (from AGC Techno Glass, or self-prepared using 10 µg/cm^2^ of rat type I collagen from Sigma-Aldrich). Passages were performed when cells were semi-confluent, by washing them with Dulbecco’s Phosphate-Buffered Saline (DPBS), followed by detachment with Trypsin-EDTA (ethylenediaminetetraacetic acid) (both from Thermo Fisher Scientific). Trypsinization was stopped by adding complete medium, and passages were performed in the range of 1:5 to 1:20 dilution. Both parental cell lines were tested free of mycoplasma contamination by Eurofins Genomics.

### Plasmid construction

The plasmids pX330 and pX459 for expression of *Streptococcus pyogenes* Cas9 and a single guide RNA were deposited to Addgene by Feng Zhang (plasmids #42230 and #48139). Target sequences were designed using the CRISPR guide RNA design tools from Benchling or CRISPOR.^52^ Selection was based on high on-target scores and low off-target scores,^2^ while avoiding targets upstream of potential alternate start codons and considering alternative splicing, using the information available in the UCSC genome browser. All the target sequences used in this study are available in supplemental table 1. Inserts were designed by adding appropriate overhangs (5’-CACC-3’ and 5’-AAAC-3’ for sense and antisense strands, respectively) to the target sequence. When the target sequence did not start with G, an additional G was added to the sense strand for the functionality of the U6 promoter, and a C was added to the end of the antisense strand make it complementary. Inserts were prepared by annealing 5 µM each of oligonucleotides (synthesized by Microsynth AG) in annealing buffer (40 mM Tris-HCl pH 8.0, 20 mM MgCl2, and 50 mM NaCl), by heating them at 95°C for 5 minutes and cooling them down slowly by reducing temperature at a speed of -0.8°C/minute in a thermal cycler. Annealed oligonucleotides (1.5 µL) were inserted by golden gate assembly into 22.5 ng of pX330 or pX459 using 0.3 µL FastDigest Bpi I (Thermo Fisher Scientific) and 0.3 µL quick ligase in 1 x quick ligase buffer (New England Biolabs) with a total volume of 6 µL. Reactions consisted of 3 cycles of incubations for 5 minutes at 37°C followed by 5 minutes at 25°C. This was followed by the addition of 0.3 µL FastDigest BpiI and 0.6 µL of water and incubation at 37°C for 1 hour to remove undigested plasmids. The plasmids pHPRTsg for expression of a single guide RNA against the rat Hprt1 or human HPRT1 gene were constructed similarly, using a previously generated pUC-U6-sg empty vector (9 ng per reaction) designed for expression of a single guide RNA under the same promoter as pX330 and pX459.^19^ Plasmids were transformed into Stbl3 chemically competent Escherichia coli (Thermo Fisher Scientific) and colonies containing correct plasmids were selected by colony PCR followed by Sanger sequencing (at Fasteris) of the PCR products (all primers used in this study are available in supplemental table 2). Plasmids were purified using the PureLink HiPure Plasmid Midiprep Kit (Thermo Fisher Scientific) or GenElute plasmid mini kit (Sigma-Aldrich). When the latter was used, plasmids were further purified by Triton X-114 isothermal extraction to obtain transfection-grade ones.^53^ Plasmids were incubated in 1% (w/v) Triton X-114 and 0.5% (w/v) sodium dodecyl sulfate (SDS) for 10 minutes at room temperature, then endotoxins were precipitated by adding sodium chloride to a final concentration of 1 M and centrifuging at 12,000 x *g* for 10 minutes at room temperature. Purified plasmids were obtained from the supernatants by isopropanol precipitation, followed by two washing steps with 70% ethanol. Plasmids were dissolved in TE buffer (10 mM Tris-HCl pH 8.0 and 1 mM EDTA).

### Generation and DNA analysis of mutant cell clones

To generate Sgpl1/Sgpp1/Sgpp2 mutant clones, three pX330-based plasmids encoding sgRNAs targeting each gene were transfected into McA-RH7777 cells using lipofectamine 3000 (Thermo Fisher Scientific). Transfected cells were seeded sparsely in 10 cm dishes to obtain colonies. When colonies grew enough to extract sufficient genome DNA for PCR, well-isolated colonies were directly lysed in lysis buffer (10 mM Tris-HCl pH 8.0, 1 mM EDTA, 0.67% SDS, and 125 µg/mL proteinase K) using cloning rings prepared by cutting pipette tips. Lysates were incubated at 55°C for 4 hours and genomic DNA was obtained by isopropanol precipitation. Mutant cells were detected by competition-based PCR using ExTaq (Takara Bio), which consists of a PCR procedure using three primers; one primer overlaps with the targeted region, while the two others flank it.^19^ Changed ratios of the amplicons was suggestive of mutations, which were further confirmed by Sanger sequencing of PCR amplicons treated with Exonuclease I and FastAP alkaline phosphatase (both from Thermo Fisher Scientific). To generate Agpat2 mutant clones, a pair of pX459-based plasmids was transfected into McA-RH7777 cells using lipofectamine 3000, followed by selection with puromycin (InvivoGen) for 24 hours starting the next day. Clones were obtained by limiting dilution in a 96 well plate. Mutant clones were selected by PCR, analyzing the deletions in genomic DNA generated by the pair of cuts.

### Generation of polyclonal mutants

McA-RH7777 cells were co-transfected with pX330-based plasmids and pHPRTsg using lipofectamine 3000. The amount of pHPRTsg used was 10% (w/w) unless stated otherwise. Seven days after transfection, 6-thioguanine (Sigma-Aldrich) was used at 4 µg/mL to select resistant cells lacking Hprt1 for 1 week. Mutant HeLa cells were generated similarly, using 1% of pHPRTsg and starting drug selection at 6 µg/mL 5 days after transfection. When POR gene was co-targeted instead of HPRT1, plasmids based on pUC-U6-sg were used to express the sgRNA targeting POR (1% of total transfected DNA). Cells were selected with 200 µM paraquat (Sigma-Aldrich) 5 days after transfection for 4 days and replated to new dishes. The efficiency of gene mutations and the indel patterns were analyzed by isolating genomic DNA as stated above, Sanger sequencing target regions amplified by PCR (supplemental tables 1 and 2), and analyzing them with TIDE (tracking indels by decomposition). When comparing the yields of resistant cells, a similar procedure was performed, and cells were fixed with methanol. Fixed cells were stained with 0.5% (w/v) crystal violet prepared in 20% methanol.

### Analysis of S1P

Cells were seeded at 750,000 cells/dish in 6 cm dishes. 24 hours later, cells were scraped in ice-cold PBS, centrifuged at 2,500 rpm, and pellets were snap frozen in liquid nitrogen. Cells were resuspended in 150 µL of pyridine extraction buffer (ethanol: water: diethyl ether: pyridine = 15: 15: 5: 1, supplemented with 2.1 mM ammonium hydroxide), to which internal standards (40 pmol C17 sphingosine, 40 pmol of C17 sphinganine, 400 pmol C17 sphingosine-1-phosphate, and 400 pmol C17 sphinganine-1-phosphate) were added. Samples were vortexed for 10 minutes at 4°C, then incubated at 4°C for 20 minutes. Samples were centrifuged at 14,000 rpm for 2 minutes at 4°C and supernatants were collected. An additional extraction was performed from remaining pellets using the same steps except for the lack of incubation. The supernatants were combined, split into two, and dried under a stream of nitrogen. Sphingoid bases were derivatized with AQC (6-aminoquinolyl-N-hydroxysuccinimidyl carbamate) by resuspending the extracts in 10 µL of 0.1% formic acid, to which was added 70 µL of borate buffer (200 mM boric acid pH 8.8, 10 mM tris(2-carboxyethyl)-phosphine, 10 mM ascorbic acid, and 33.7 mM ^15^N^13^C-valine; the heavy isotope-labeled valine was included for potential analysis of amino acids, not performed in this study) and 20 µL of 2.85 mg/mL AQC in acetonitrile. Derivatization was performed for 15 minutes at 55°C and overnight at 24°C. Samples were centrifuged at 14,000 rpm for 2 minutes, and supernatants were analyzed by LC-MS/MS in an Accela HPLC system coupled to a TSQ Vantage mass spectrometer (Thermo Fisher Scientific). Samples were separated on a C18 column (EC 100/2 Nucleoshell RP-18, 2.7 µm, from Macherey-Nagel) using a gradient of 0.1% formic acid / isopropanol over 0.1% formic acid / water. Multiple reaction monitoring was used to detect AQC-derivatized S1P at Q1 (*m/z* = 536.3 for C17 S1P and 550.3 for C18 S1P) and the AQC fragment at Q3 (*m/z* = 171.1). Concentrations were calculated by normalizing signals for extraction efficiency using signals of the C17 internal standard, which was further corrected for signal responses that were measured by injecting synthetic standards in another run. To normalize variations in cell numbers between samples, pellets generated during sample extractions were washed with ethanol, dried for 5 minutes at room temperature, and lysed in protein lysis buffer (100 mM Tris-HCl pH 8.0, 3 M Urea, and 1% SDS). Samples were incubated at 60°C for 15 minutes and sonicated for 5 minutes in a bath sonicator. The incubation-sonication cycle was repeated once more. Protein concentrations were analyzed with a Bicinchoninic Acid Protein Assay Kit (Sigma-Aldrich), using bovine serum albumin as a standard, which were used to normalize variations in cell numbers.

### Analysis of phosphatidylcholine (PC) by LC-MS/MS

McA-RH7777 cells were seeded on collagen-coated 96 well plates at 10,000 cells/well. When analyzing polyunsaturated fatty acid (PUFA)-supplemented cells, 10 µM each of linoleic acid, arachidonic acid, and docosahexaenoic acid was added the next day. 24 hours later, cells were washed with PBS and methanol was added to extract PC and incubated at 4°C for >1 hour. Samples were centrifuged at 13000 rpm for 5 minutes to remove debris, and the supernatants were analyzed by LC-MS/MS in an ACQUITY UPLC system coupled to a TSQ Vantage mass spectrometer. Samples were separated on a C8 column (ACQUITY UPLC BEH C8, 1 x 100 mm, 1.7 µm, Waters) using a gradient of acetonitrile over 20 mM ammonium bicarbonate. We noted severe peak tailing of some PC species depending on the condition of the LC-MS/MS system, and in such cases acetonitrile was replaced by isopropanol: acetonitrile = 3: 7. PC was detected by a precursor ion scan detecting the phosphocholine fragment of *m/z* = 184.1 at Q3. Since the biological question was to analyze the acyl-chain balance of PC, the sum of PC signals was used for normalization.

### Analysis of fatty acids by GC

McA-RH7777 cells were seeded on 12 well collagen-coated plates at 120,000 cells/well and PUFAs were supplemented the next day as described above. 24 hours later, cells were washed with PBS, and scraped in methanol. Methanol was further added to the wells to maximize recovery and combined with the scraped cells. Lipids were extracted by the method of Bligh and Dyer, the extracts were resuspended in chloroform, and were loaded on InertSep NH2 aminopropyl columns (GL Science). Neutral lipids were eluted with chloroform: isopropanol = 2: 1, free fatty acids were eluted with 2% acetic acid in diethyl ether, and phospholipid were collected by eluting them in 2.8% ammonia in methanol. C23:0 (Supelco n-Tricosanic acid) was added as a standard to the phospholipid fractions. Fatty acid methyl esters were prepared with the Fatty Acid Methylation Kit, and purified using the Fatty Acid Methyl Ester Purification Kit (both from Nacalai Tesque). Fatty acid methyl esters were separated on a capillary column (FAMEWAX, 30 m, 0.25 mm ID, 0.25 µm from Restek) using a GC-2010 Plus system equipped with a flame ionization detector (Shimadzu). Sums of abundant fatty acids were used to normalize the data and calculate the percentages of individual fatty acids. See reference for detailed procedures.^54^

### Lipidome analysis by nanoESI-MS

Cells were seeded on 6 cm dishes at 750,000 cells/dish. 24 hours later, cells were scraped in ice-cold PBS, pelleted down by centrifugation at 2,500 rpm at 4°C, and pellets were snap frozen in liquid nitrogen. Lipids were extracted using a modified methyl tert-butyl ether (MTBE) extraction protocol. Pellets were resuspended in 200 µL 50% methanol/water, 100 µL of an internal standard mix (400 pmol dilauryl-PC, 1 nmol 17:0-14:1 phosphatidylethanolamine, 1 nmol 17:0-14:1 phosphatidylinositol, 3.3 nmol 17:0-14:1 phosphatidylserine, 700 pmol tetramyristoyl cardiolipin, 2.5 nmol C17 ceramide, 500 pmol C8 glucosylceramide, and 100 pmol C12 sphingomyelin, resuspended in chloroform: methanol = 1: 1 and diluted in methanol, all from Avanti Polar Lipids) was added, followed by 180 µL methanol and 1.2 mL MTBE. Samples were vortexed for 10 minutes at 4°C and incubated at room temperature for 1 hour. 200 µL of water was added to induce phase separation. After 10 minutes of incubation, samples were centrifuged at 10,000 x *g* for 10 minutes at room temperature. The upper layers were transferred to glass tubes, and the remaining samples were used for a re-extraction with 400 µL of artificial upper layer (MTBE: methanol: water = 10: 3: 1.5), which was incubated and centrifuged as described above. Upper layers were combined (designated total lipid fractions), split into three vials, and dried under a nitrogen stream. Among the split samples, one was further treated by a mild base to remove glycerophospholipids, which was critical to reduce ion suppression of hexosylceramide species and to detect sphingomyelin accurately without interference from isotopes of PC. For this, dried samples were reconstituted in 500 µL of methylamine solvent (methanol: water: water-saturated butanol: methylamine = 4: 3: 1: 5), sonicated for 5 minutes in a bath sonicator, and incubated for 1 hour at 53°C. Samples were dried under a nitrogen flux, followed by desalting. For desalting, dried samples were reconstituted in 300 µL water-saturated butanol, sonicated for 5 minutes in a bath sonicator, and 150 µL water was added to induce phase separation. Samples were centrifuged at 3200 x *g* for 10 minutes at room temperature, and upper layers were transferred to glass vials. Another extraction was repeated by adding 300 µL water-saturated butanol to the remaining samples and combined with the first extracts. Samples were dried under a nitrogen flux and were designated as sphingolipid fractions. Lipids from each fraction were reconstituted in 250 µL chloroform: methanol = 1: 1 and diluted 4 and 10 fold for samples to be measured in the negative and positive ion mode, respectively. Dilutions were done in chloroform: methanol: water = 2: 7: 1 for samples analyzed in the positive ion mode and chloroform: methanol = 1: 2 for samples analyzed in the negative ion mode, both being supplemented with 5 mM ammonium acetate. Diluted samples were directly infused in a TSQ Vantage mass spectrometer using a robotic sample delivery system equipped with a nanoESI source (TriVersa NanoMate from Advion). Lipids were detected by multiple reaction monitoring, using fragments specific for each lipid classes. A full list of transitions is available in supplemental table 3. Individual lipids were quantified by dividing signal intensities by those of corresponding internal standards and then multiplying them with the amount of internal standard added. Then sums of each lipid class were calculated after removing low abundance lipids. For this filtering, the averages of the five lowest values for each lipid class were used to estimate the noise levels, and only signals at least two-fold higher than this noise were considered. Due to the low numbers of sphingolipids and sterols analyzed, this filtering was performed only for glycerophospholipids. The sums of all major lipids were calculated from phosphatidylcholine, ether phosphatidylcholine, phosphatidylethanolamine, ether phosphatidylethanolamine, phosphatidylinositol, phosphatidylserine, cardiolipin, ceramide, hexosylceramide, and sphingomyelin. Lysophospholipids and sterols were not used to calculate the sums. The sums were used to normalize data variances coming from the small differences in sample amounts used for lipid extraction, and thus lipid data are illustrated as “% of total”. This normalization was used empirically as an efficient way to generate consistent datasets between replicates, making the detection of differentially abundant lipids more robust.

### Untargeted lipidomics by LC-MS

Cells were seeded on 6 cm dishes at 750,000 cells/dish. 24 hours later, cells were scraped in ice-cold PBS, pelleted down by centrifugation at 2,500 rpm at 4°C, and pellets were snap frozen in liquid nitrogen. Total lipids were extracted by the method of Bligh and Dyer. Pellets were resuspended on ice in 200 µL water, 500 µL of methanol was added, followed by addition of internal standards (4 µL of SPLASH LIPIDOMIX Mass Spec Standard and 20 pmol each of C17 ceramide, C17 Glucosylceramide, and C17 Lactosylceramide, all from Avanti Polar Lipids). Then 250 µL of chloroform was added and the single phase was shaken for 10 minutes. Phase separation was initiated by the sequential addition of 250 µL of water and 250 µL of chloroform, and the mixture was shaken for 10 minutes. Tubes were centrifuged for 15 minutes at 3,000 rpm. 400 µL of lower phase were transferred to new glass tubes and evaporated under a stream of nitrogen. The extracts were resuspended in 60 µL isopropanol: methanol: water (5:3:2) and 3 µL was injected into the LC-MS, consisting of a Ultimate 3000 UHPLC system coupled to a Q Exactive Plus mass spectrometer (Thermo Fisher Scientific). Lipids were separated on an Accucore C18 column (150 x 2.1 mm, 2.6 µm particles), using a gradient of solvent B (isopropanol: acetonitrile: water (88:10:2, v/v) supplemented with 2 mM ammonium formate and 0.02% formic acid) over solvent A (acetonitrile: water (1:1, v/v) supplemented with 10 mM ammonium formate and 0.1% formic acid) at a flow rate of 400 µL/minute. The gradient started at 35% solvent B and was changed linearly to reach 60% B at 4 minutes, 70% B at 8 minutes, 85% B at 16 minutes, 97% B at 25 minutes, and 100% B at 25.1 minutes. 100% B was maintained until 31 minutes, and the column was reconditioned at 35% B for 4 minutes. Separate injections were done to analyze lipids in positive and negative ion modes. Data were acquired in a data dependent MS2 mode. MS1 was analyzed at a resolution of 70,000 at *m/z* = 200 on the range of *m/z* = 250-1200 with an AGC target of 1000000 and maximum injection time of 250 msec. For MS2, 15 precursor ions were selected in an isolation window of *m/z* = 0.95 and fragmented by higher-energy collisional dissociation with a normalized collision energy of 25 eV and 30 eV for positive mode and 20 eV, 30 eV, and 40 eV for negative mode, and injected into the Orbitrap analyzer with an AGC target of 100000 and maximum injection time of 80 msec, and analyzed at a resolution of 35,000 at *m/z* = 200. Lipids were identified by their fragmentation patterns using MS-DIAL5 ^12^. Data was manually curated based on retention times and investigation of co-eluting peaks, after which peak areas were collected. Lipids were semi-quantified by dividing peak areas with those of corresponding internal standards and multiplying with the quantities that were added to samples. Lipid levels were normalized with the sum of ceramide, hexosylceramide, dihexosylceramide, trihexosylceramide, sphingomyelin, lysophosphatidylcholine, phosphatidylcholine, lysophosphatidylethanolamine, phosphatidylethanolamine, phosphatidylinositol, and phosphatidylserine.

### Measurement of lysophosphatidic acid acyltransferase (LPAAT) activity

Cells were disrupted on ice using a probe sonicator (Ohtake Works) through 2 rounds of sonications for 30 seconds each in TSC buffer (100 mM Tris-HCl pH 7.4, 300 mM sucrose, and 1 x complete protease inhibitor cocktail). Sonicated samples were centrifuged at 800 x *g* for 10 minutes, and supernatants were centrifuged at 100,000 x *g* for 1 hour to pellet membrane fractions. Pellets were resuspended in TSE buffer (20 mM Tris-HCl pH 7.4, 300 mM sucrose, and 1 mM EDTA) and protein concentrations were measured using the Protein Assay Kit (Bio-Rad). LPAAT activity was measured by incubating membrane fractions (containing 0.01 µg of protein) with substrates at 37°C for 10 minutes. Reaction mixtures contained 25 µM deuterium-labeled lysophosphatidic acid, 1 µM each of 16:0-, 18:1-, 18:2-, 20:4-, and 22:6-CoA (Avanti Polar Lipids), 110 mM Tris-HCl pH 7.4, 150 mM sucrose, and 0.5 mM EDTA. Reactions were stopped by the addition of chloroform: methanol = 1: 2. Dimyristoyl phosphatidic acid (Avanti Polar Lipids) was added as an internal standard, and Bligh and Dyer extraction was performed. Reaction products were quantified using LC-MS/MS by comparing peak areas of chromatograms to those of lipid standards. The system was the same as the one used for PC analysis, but the column was an ACQUITY UPLC BEH amide column (2.1 x 30 mm, 1.7 µm, from Waters). See reference for detailed procedures.^33^

### Informatics and Data analysis

Gene essentiality across multiple cell lines was analyzed using the depmap portal website, where data of genome-wide CRISPR screenings in 789 cell lines is available.^50^ Raw data were downloaded from the website and values were extracted with R software. Gene expression profiles of Agpat genes across multiple mouse tissues were obtained from BioGPS.^51^ LPAAT activity in multiple tissues was analyzed previously.^33^ The correlation between datasets was calculated from the data of lung, heart, liver, spleen, kidney, stomach, small intestine, colon, testis, adipose tissue, and skeletal muscle, which are the ones overlapping between the two. Statistical analyses were performed using Prism 8 (GraphPad). Multivariate analyses (principal component analysis, K-means clustering, and hierarchical clustering) were performed using R.

---

**Figure S1.**
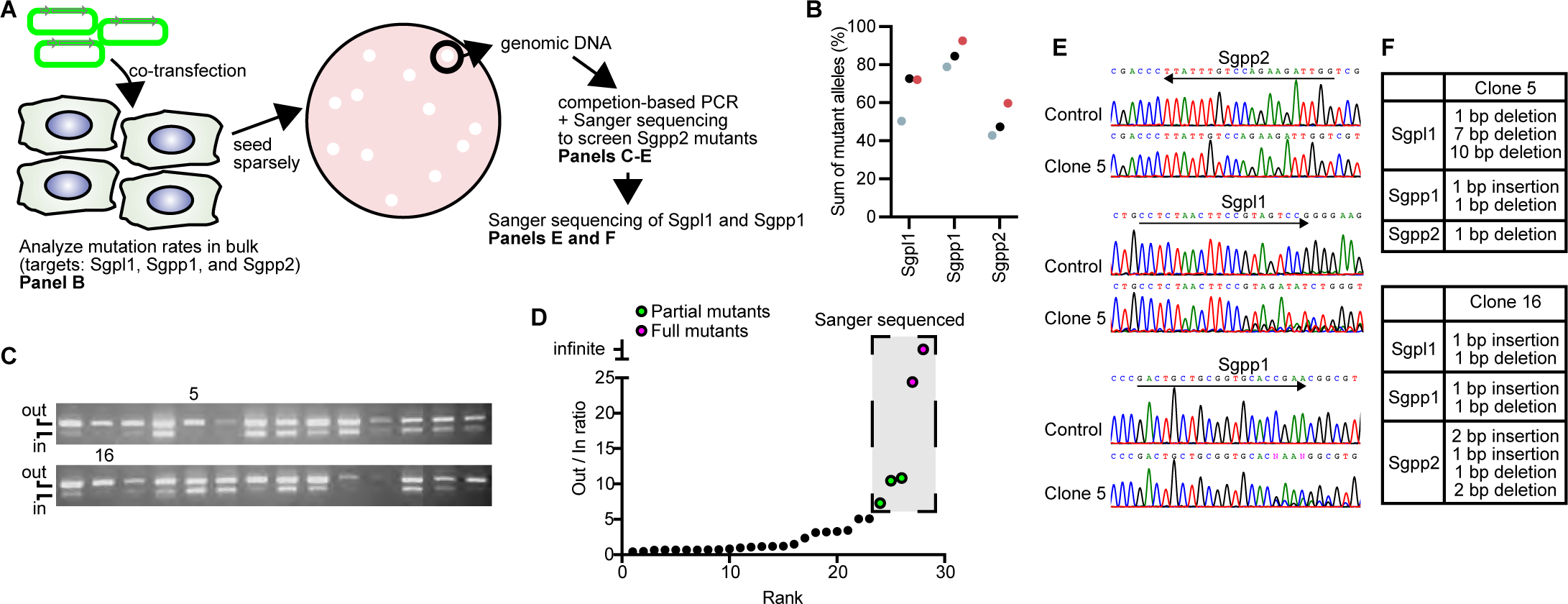
Preliminary experiment leading to the idea of GENF. (A) Overview of the experiment. McA-RH7777 cells were co-transfected with three pX330-based plasmids encoding Cas9 and sgRNAs against Sgpl1, Sgpp1, and Sgpp2. Transfected cells were analyzed in bulk to test the efficiencies of different sgRNAs, while one part was reseeded in dishes sparsely to obtain colonies. Colonies were directly lysed to obtain genomic DNA and screen for Sgpp2 mutants. Sgpl1 and Sgpp1 mutation was analyzed in the obtained Sgpp2 mutants. (B) In three experiments (distinguished by colors), mutation rates of Sgpp2 were always the lowest after transiently co-transfecting pX330-based plasmids. (C and D) Screening of Sgpp2 mutants by competition-based PCR. The method uses a mixture of primer that enables the detection of mutant clones by a reduction of the amplicon labeled “in” compared to the one labeled “out”, since one of the primers for “in” amplicon binds to the sgRNA target. See reference^19^ for details. (C) Result of competition-based PCR. PCR patterns in the numbered lanes are from the clones that were later confirmed to be fully mutated. (D) Quantification of signal ratios between the two amplicons in (C). Five clones were selected for Sanger sequencing, and two of them were full mutants. (E) Sanger sequencing of the three targets in parental cells or clone 5. Target regions are underlined with an arrow. (F) Summary of mutations seen in clones 5 and 16. Note the absence of wild type alleles in Sgpl1 and Sgpp1, despite the fact that the clones were pre-selected based only on Sgpp2 mutations.

**Figure S2.**
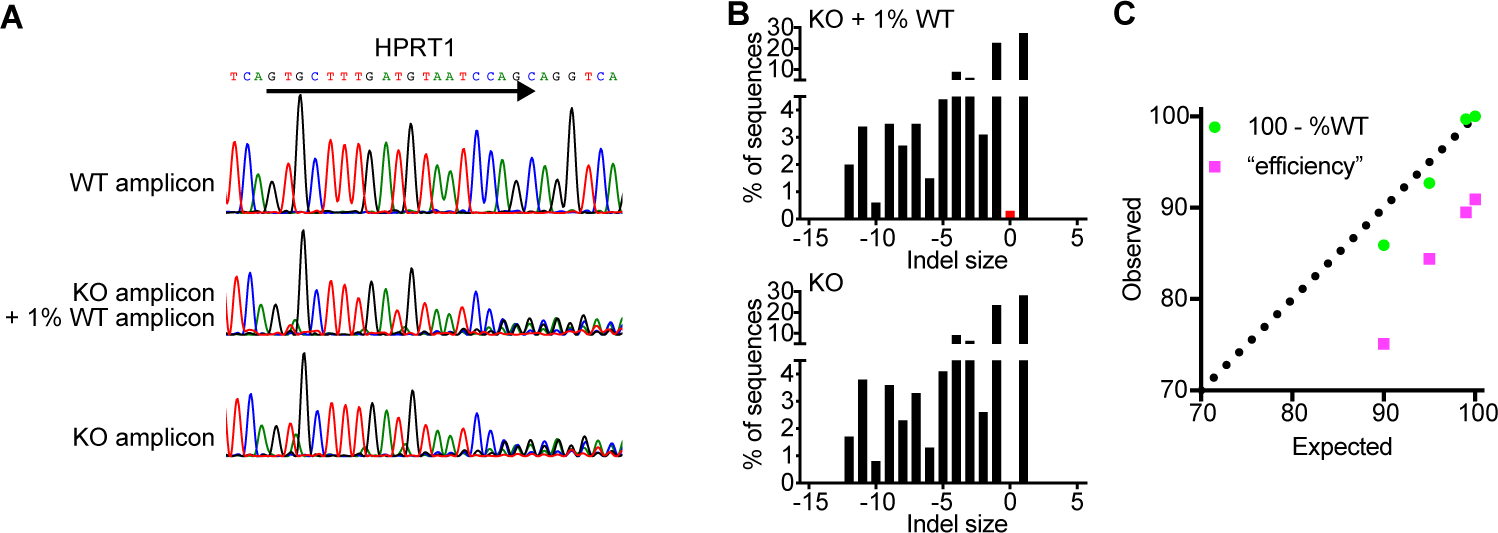
Evaluation of a method to analyze high mutation rates. (A) Sanger sequencing of PCR amplicons from wild type or mutant HPRT1 locus, either alone or mixed at the indicated ratio. (B) TIDE (tracking indels by decomposition) analysis^21^ from the Sanger sequencing results of (A). The calculated frequency of alleles with the indicated indel size is illustrated. Note that as little as 1% of wild type allele can be detected (with an indel size of zero, red bar in top panel) with TIDE. (C) Comparison of expected mutation rates (from the ratio of mixed PCR amplicons) and observed mutation rates calculated in two ways. Observed mutation rates were calculated either from the sum of mutant alleles detected with TIDE (“efficiency”, which is usually used) or by subtracting the detected wild type alleles from 100 (“100-%WT”). Note that the latter gave a better estimate of mutation rates.

**Figure S3.**
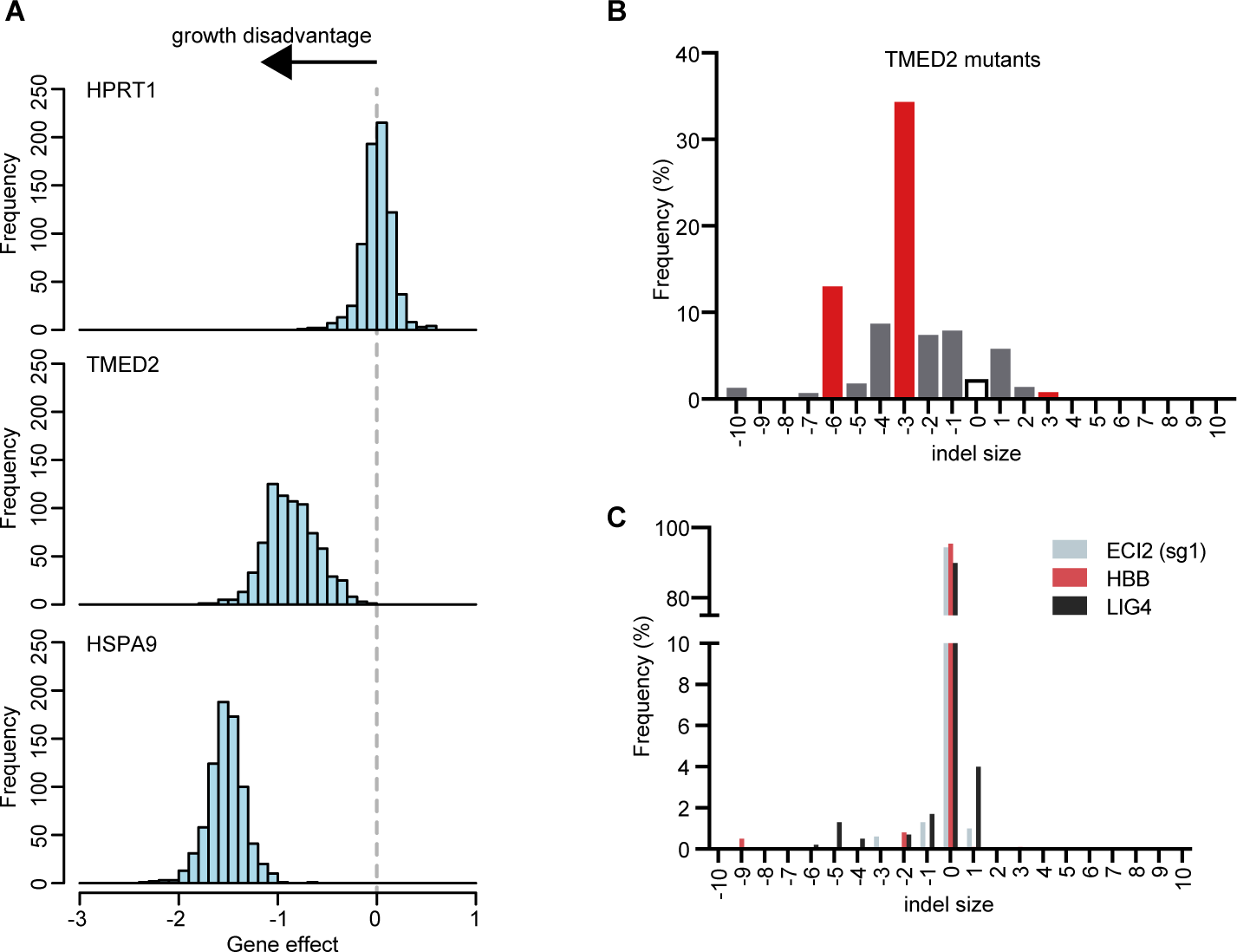
Analysis of failed cases of GENF. (A) Essentiality of selected targets. Gene effect scores were obtained from Depmap database.^50^ The scores illustrate the degree of sgRNA depletion/enrichment from cell populations during genome-wide CRISPR-Cas9 screenings in multiple cancer cell lines. A shift of the traces to the left (as seen for HSPA9 and TMED2) illustrate that the sgRNA targeting the gene is depleted upon culture, showing that the target is critical for cell survival in multiple cell lines. Note that HPRT1, which was used for co-targeting throughout this study, is not essential. (B) Indel patterns in TMED2 mutants analyzed by TIDE. Note that this target could be highly mutated with GENF with barely detectable wild type alleles (white bar), but with a high enrichment of in-frame mutations (red bars). (C) TIDE analysis of cells transiently transfected with pX330-based plasmids encoding the sgRNAs targeting the indicated genes. Note the very low levels of mutation induced with the sgRNAs targeting ECI2 or HBB. The result of a similar transient transfection to mutate LIG4 is illustrated to compare the efficiency, which shows that a slightly better target sgRNA is sufficient to obtain nearly complete gene disruption with GENF (see LIG4 mutation rates with GENF in Figure 1J).

**Figure S4.**
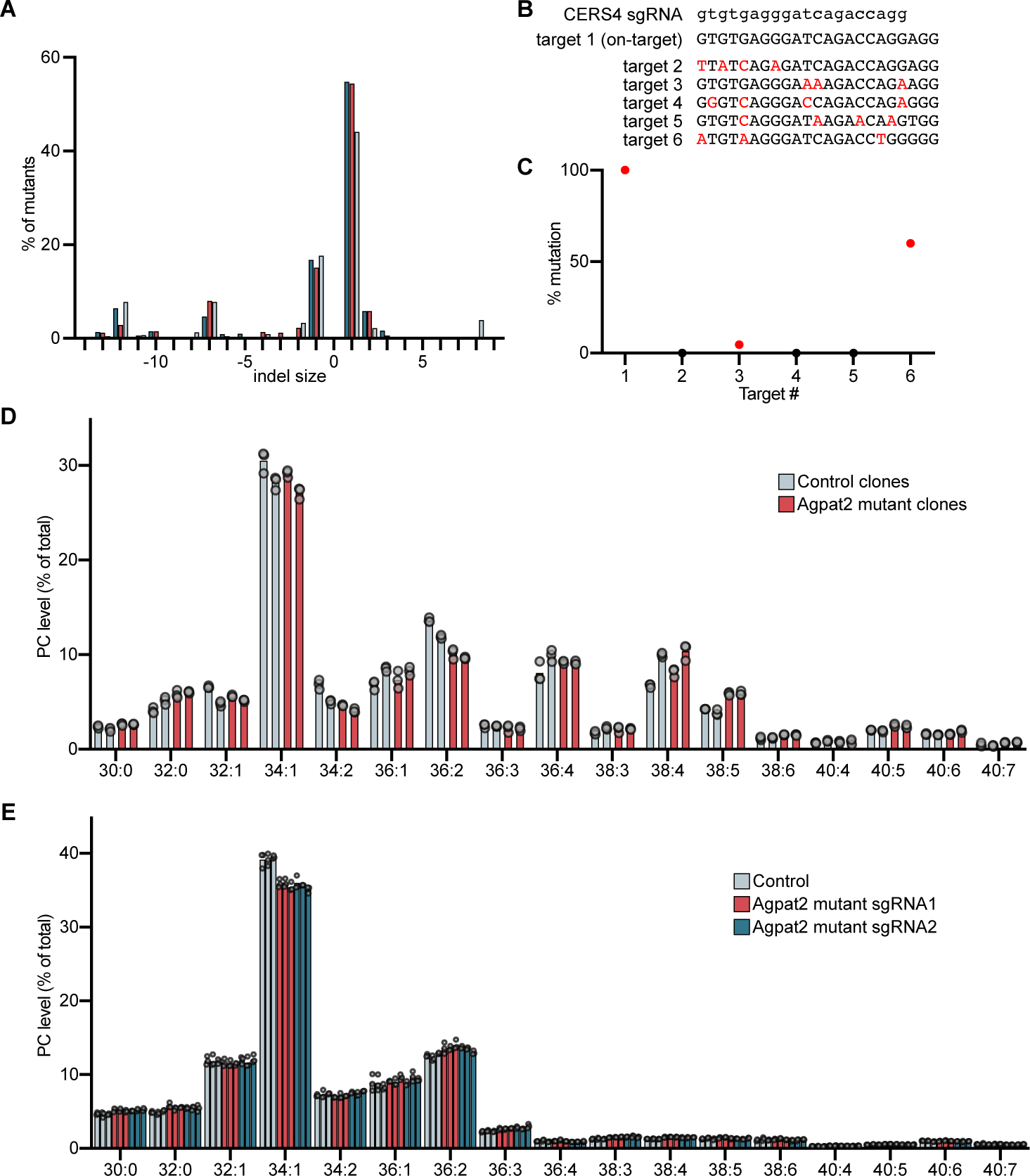
Additional data to characterize GENF. (A) Indel patterns of CERS6 mutants generated on different days, shown in different colors. (B) Off-target sites of CERS4 sgRNA predicted based on homology, with mismatches highlighted in red. (C) On- and Off-target mutations observed in CERS4 mutant cells. Non-zero values are plotted in red. (D) Comparison of PC composition between control and Agpat2 mutant clones. Different clones are shown as distinct bars. (E) Comparison of PC composition between polyclonal control and Agpat2 mutant cells generated with GENF. Different polyclonal cell lines are shown as distinct bars. We know empirically that the content of polyunsaturated glycerophospholipids vary largely between datasets obtained on different days, which is likely due to differences in various factors including the conditions of the serum, which is the sole source of polyunsaturated fatty acids in standard culture conditions. The differences in control cells between (D) and (E) are likely to be caused by this variability, since the datasets were obtained years apart with different batches of sera.

**Figure S5.**
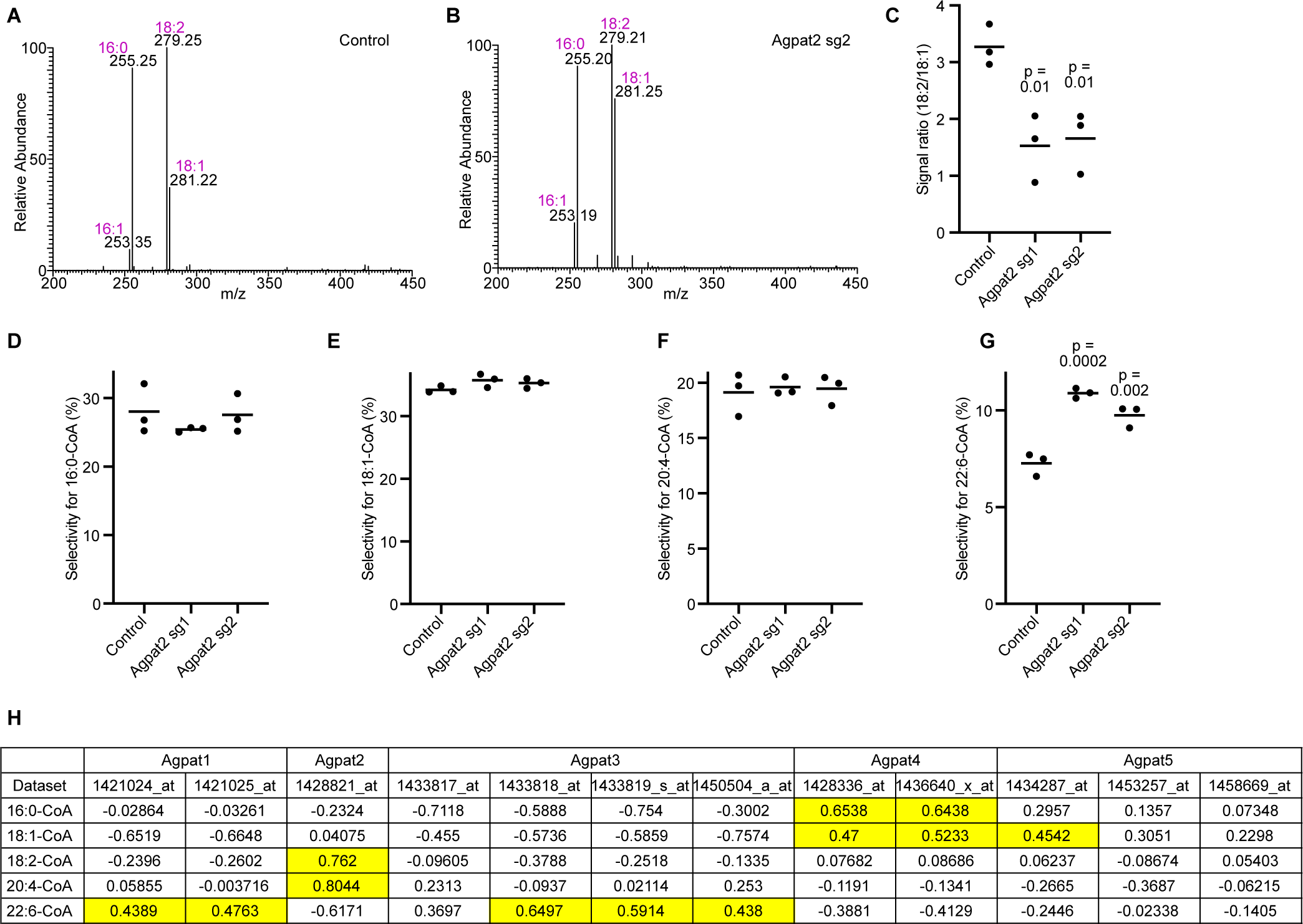
Agpat2 as a regulator of linoleic acid levels. (A-C) Fragmentation patterns of PC 34:2 analyzed by tandem mass spectrometry. PC 34:2 from control (A) and Agpat2 mutant (B) cells was analyzed by liquid chromatography-tandem mass spectrometry as a bicarbonate adduct negative ion and fragmented to reveal the fatty acyl composition. Fragments corresponding to fatty acids are annotated. Similar results were obtained in three biological replicates and are quantified in (C). (D-G) Selectivity of lysophosphatidic acid acyltransferase (LPAAT) activity in membrane fractions obtained from control or mutant cells for the indicated substrates. (C-G) p values: one-way ANOVA followed by Dunnett’s multiple comparisons test to compare with Control. (H) The correlation between LPAAT acyl-CoA selectivity^33^ and mRNA expression^51^ of various LPAAT enzymes in various tissue. Values above 0.4 are colored.

**Figure S6.**
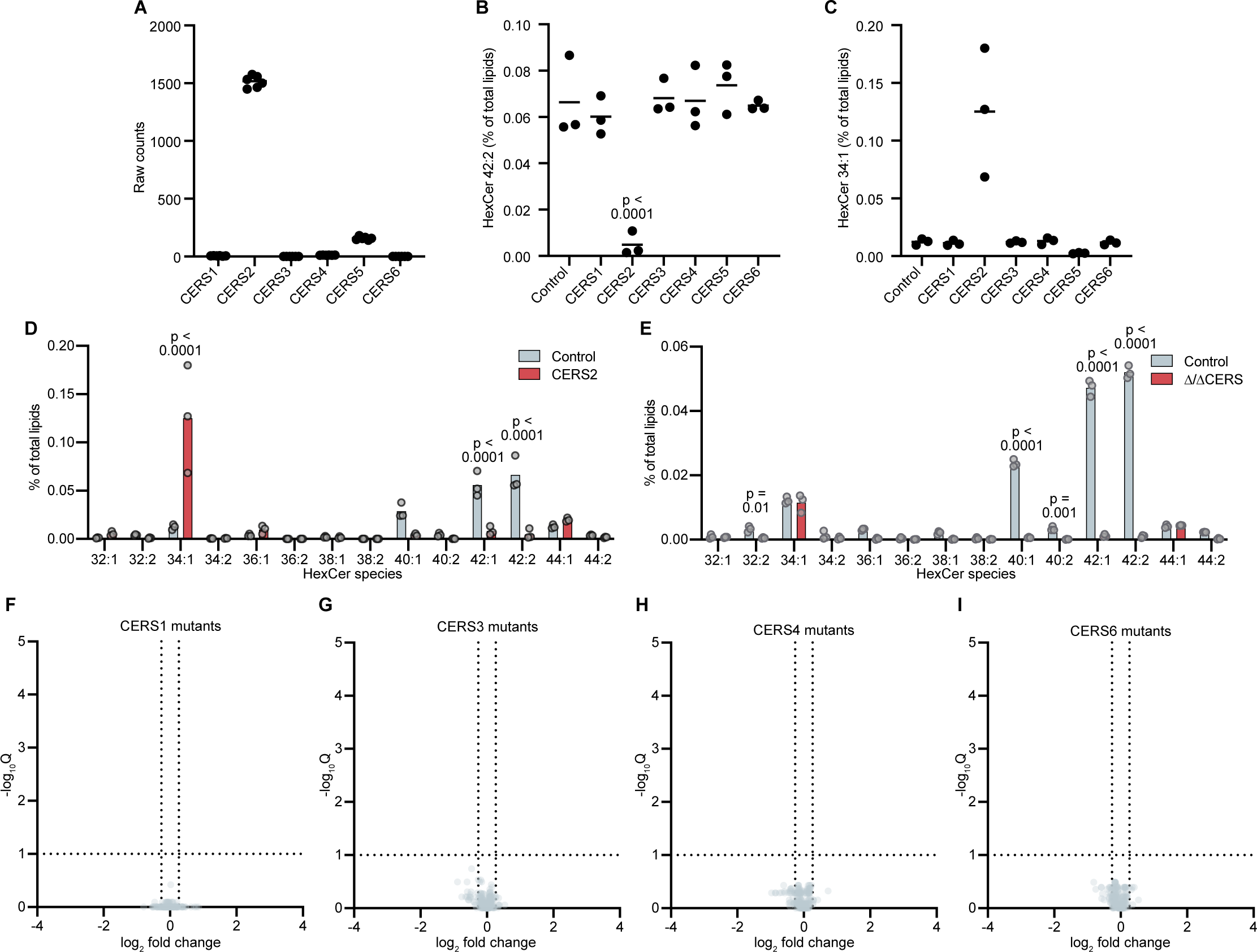
Lipid changes in CERS mutants. (A) Expression of CERSs in HeLa cells obtained from RNA-seq ^39^. (B and C) Changes in the indicated Hexosylceramide (HexCer) species in CERS mutants. (B) p values: one-way ANOVA followed by Dunnett’s multiple comparisons test to compare with Control. (D and E) Changes in HexCer species in CERS2 and Δ/ΔCERS mutants. p values: two-way ANOVA followed by Sidak’s multiple comparisons test. (F-I) Volcano plots illustrating changes in lipid levels and statistical their significances seen in the indicated CERS mutants. No lipid is identified as a hit (having changes above 1.2-fold and q values (FDR-corrected multiple t-test) below 0.1).

